# Protein import-driven mitochondrial hyperactivation dictates angiogenic signalling independently of HIF-1α

**DOI:** 10.64898/2026.04.11.717880

**Authors:** Tanvi Chaudhary, Sharath Mohan Bhat, Priyadarshika Pradhan, Manjunath B Joshi, Devanjan Sinha

**Author notes:** To whom correspondence should be addressed: Dr. Devanjan Sinha, Department of Zoology, Institute of Science, Banaras Hindu University, Varanasi, 221005, India.

## Abstract

Angiogenesis is essential for sustained neoplastic progression, yet its initiation has largely been framed through hypoxia-dependent HIF signalling. In OXPHOS driven tumours, how mitochondrial function contributes to angiogenic induction remains poorly understood. Here, we uncover a non-canonical mechanism linking tumour cell mitochondrial hyperactivity to a pro-angiogenic response. Mitochondrial function critically depends on the import of nuclear-encoded proteins, largely mediated by TIM23 translocase. We found Timm44, the central scaffold of this translocase, to be broadly upregulated across angiogenesis-dependent malignancies. Using gain-of-function system that recapitulate tumour-associated Timm44 amplification, we enhanced mitochondrial protein import capacity, thereby remodelling mitochondrial gene expression and driving hyperactivation of the electron transport chain, particularly Complex I. This hyperactive mitochondrial state generated a signalling-competent ROS-enrich environment that induces VEGFA expression independently of HIF-1α stabilization, via a redox-licensed TRX–ASK1–p38MAPK–SP1 axis, thereby promoting endothelial sprouting and proliferation. These findings define protein import–driven mitochondrial hyperfunction as an alternative driver of tumour angiogenesis, extending current paradigms beyond classical hypoxia-dependent mechanisms and highlighting new therapeutic vulnerabilities.

## Introduction

Angiogenesis, the ability of endothelial cells (ECs) to sprout new capillaries from pre-existing vessels is a defining hallmark of an aggressively progressing neoplasm(*1–3*). By releasing pro-angiogenic factors such as VEGFs, cancer cells recruit ECs to proliferate, migrate, and assemble into new vascular networks that ultimately supply the expanding tumor with oxygen and nutrients(*4–6*). This vascular expansion not only sustains tumor growth but also creates direct routes for metastasis and shapes the tumor’s immune landscape(*7, 8*). Although classical signalling pathways involving VEGFA, FGF, and PDGF have long been recognized as principal drivers of angiogenesis(*9–11*), it has become increasingly clear that these pathways do not function in isolation. Instead, they are profoundly shaped by the metabolic environment in which they operate.

Under normal physiological conditions, angiogenesis is tightly regulated by a balance of pro-and anti-angiogenic cues that preserve vascular stability(*12*). In tumors, this balance collapses as proliferating cancer cells quickly outpace their blood supply, generating regions of severe hypoxia. The cellular response to hypoxia hinges on the stabilization of hypoxia-inducible factors (HIFs), which activate pro-angiogenic transcriptional programs, including VEGFA(*13*). Even under normoxic conditions, mutations in the oxygen-sensing protein VHL can stabilize HIF-1α, illustrating how deeply tumors rely on this pathway to maintain their angiogenic drive(*14*). Early models of tumor angiogenesis emerged largely from glycolytic tumors, where inhibition of PHD2 by lactate accumulation or redox imbalance, along with inflammation, fuel HIF-dependent angiogenic response(*15–17*).

Yet this classical view captures only part of the story. Many solid tumors do not operate purely as glycolytic systems. Instead, they retain active, adaptable mitochondria and rely on oxidative phosphorylation (OXPHOS), TCA cycle flux and anaplerotic substrates to meet the biosynthetic and energetic demands of rapid proliferation(*18*). Mitochondrial adaptability is increasingly recognized as a key determinant of tumor fitness, supporting their growth and metastatic dissemination by dynamically rewiring OXPHOS and metabolite-driven signaling pathways(*19–21*). They help maintain redox balance, buffer metabolic stress, and influence how tumors respond to therapy.

Insights into the metabolic control of angiogenesis have largely emerged from endothelial biology, where ECs exhibit remarkable metabolic plasticity(*22*). ECs switch between glycolysis, fatty acid oxidation, glutamine metabolism, and OXPHOS depending on their role during vessel sprouting, migration, or stabilization(*23–25*). In contrast, how tumor-cell mitochondria, and not endothelial mitochondria, influence angiogenesis is far less understood, despite the fact that many tumors depend heavily on OXPHOS-dependent states. How such tumors achieve high angiogenic capacity without relying solely on classical hypoxia-driven HIF pathways remains unresolved.

A major determinant of mitochondrial performance is the import of nuclear-encoded proteins through the TOM and TIM complexes because OXPHOS-dependent tumors must maintain high mitochondrial protein import to sustain respiratory chain activity and metabolic output. Of these, TIM23-complex serves as the principal conduit for import of matrix-targeted proteins(*26–28*). Timm44, a central scaffold of this complex, orchestrates the recruitment of mtHsp70 and its cofactors to drive ATP-dependent protein translocation(*29, 30*). Timm44 levels rise under metabolic or oxidative stress, boosting mitochondrial resilience, whereas its depletion disrupts proteostasis and compromises cell viability across multiple model systems, including ECs(*31–34*). Despite its clear importance for mitochondrial health, the contribution of Timm44 to tumor metabolism and angiogenesis has remained underexplored.

Given its central role in mitochondrial proteostasis, we investigated whether Timm44 could influence how tumor cells regulate angiogenesis. We found that Timm44 is broadly overexpressed in angiogenesis-driven tumors. Functional elevation of Timm44 enhanced mitochondrial protein import efficiency, expanded mitochondrial DNA content and translational capacity, and shifted tumors toward greater OXPHOS dependence through augmented Complex I and III subunit levels. This respiratory upregulation activated redox-sensitive transcription factors such as SP1, ultimately driving HIF1α-independent VEGFA induction. By integrating TCGA-dataset analyses with molecular and functional validation, our study identifies Timm44-driven mitochondrial proteostasis as a previously unrecognized regulatory axis through which tumor cells orchestrate angiogenic signaling. These findings highlight a new dimension of mitochondrial control over tumor vascular adaptation and suggest that Timm44 may represent a metabolic lever through which tumors sustain their angiogenic state.

## Results

### Angiogenesis driven malignancies show elevated Timm44 expression

Mitochondria function not only as bioenergetic centers but also as dynamic hubs that make the tumors stress resilient and help them to sustain unchecked growth(*2, 35*). Because mitochondrial performance depends heavily on the efficient import of nuclear-encoded proteins, we examined the key element of this machinery, the TIM23-complex, specifically its core scaffold protein Timm44 which coordinates recruitment of mtHsp70, guiding nuclear-encoded proteins into the matrix(*27, 36*).

Timm44 was found to be consistently elevated across multiple tumor types. Integrated analysis across pan-cancer transcriptomic datasets (TCGA and GTEx) using UALCAN, TIMER2, and UCSC Xena platforms revealed upregulation of Timm44 transcripts in several cancer cohorts relative to their normal counterparts. This enrichment was particularly pronounced in angiogenesis-driven malignancies, including kidney renal clear cell carcinoma, lung adenocarcinoma, colon adenocarcinoma, Breast cancer invasive carcinoma and cholangiocarcinoma tumors (Fig. S1A). Supporting this observation at the protein level, immunohistochemistry from the Human Protein Atlas revealed strong Timm44 staining in malignant epithelial compartments, whereas adjacent normal parenchyma displayed only faint expression (Fig. S1B). Enhanced expression of Timm44 was observed across primary, recurrent, and metastatic tumors in the TCGA pan-cancer cohort (Fig. S1C). Importantly, Timm44 remained elevated across all tumor grades and stages across different subtypes, reflecting its involvement throughout disease development (Fig. S1D-G). Further analysis of the TCGA-BRCA cohort revealed that Timm44 expression was highest in primary solid tumors which typically show high angiogenic dependency (Fig. S1H), and elevation of the protein levels were found across molecular subclasses (Fig. S1I). Among BRCA cohort, patients with higher Timm44 expression experienced poorer overall survival, aligning Timm44 levels with unfavorable clinical outcomes (Fig. S1J).

To uncover what drives this overexpression, we examined Timm44 across genomic copy-number states. A clear gene-dosage effect became evident: tumors with diploid or gain states already exhibited elevated transcripts, while those harboring high-level amplifications showed the strongest upregulation (Fig. S1K). This relationship suggests that copy-number amplification is a major contributor to Timm44 overexpression in cancer. Taken together, these findings pointed to Timm44 as a potentially important mitochondrial regulator of tumor aggressiveness. Because its mRNA levels were highest in gain- and amplification-bearing tumors, we modeled this clinically relevant state using a Timm44 overexpression system in breast cancer cells. This approach allowed us to directly test how enhanced mitochondrial import capacity might influence the angiogenic program under the parallel metabolic state that fuel rapidly proliferating cancer cells.

### Timm44 primes tumor cells for angiogenic output

To determine whether the elevated levels of Timm44 observed in tumors translate into a functional angiogenic output, we overexpressed Timm44 (*Timm44^OE^*) in three breast cancer cell lines, MCF7 (Fig. S1L), T47D and ZR75 and evaluated for changes in VEGFA levels, a canonical and rate-limiting pro-angiogenic factor. Across all models, Timm44 overexpression led to a marked increase in VEGFA protein abundance (Fig. 1A-B, S2A-B), accompanied by parallel elevation in VEGFA transcript levels (Fig. 1C), suggesting a transcriptionally driven enhancement of its expression. A positive correlation was also observed between Timm44 and VEGFA in cBioPortal datasets (Fig. S2C)

**Fig 1.**
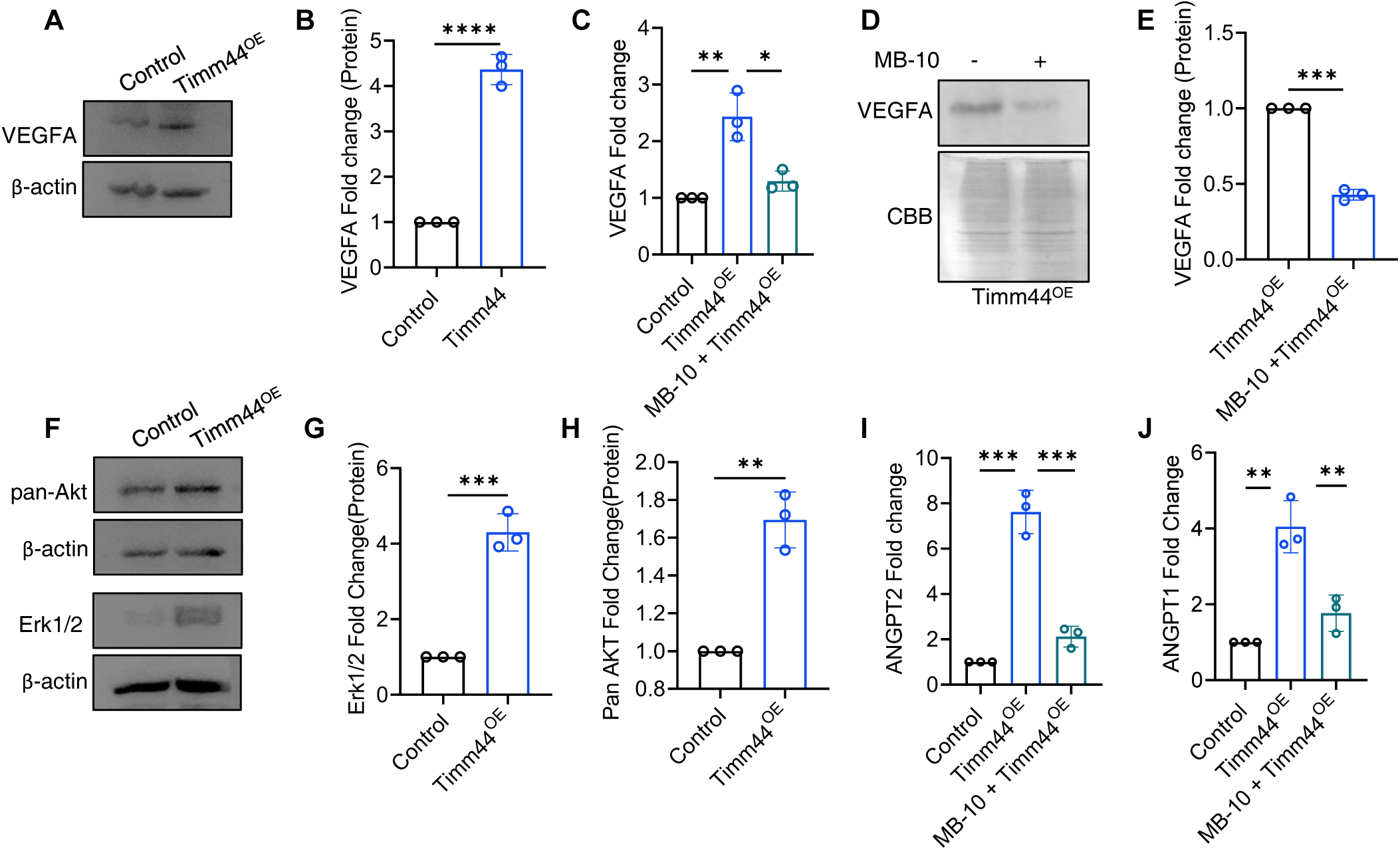
Timm44 expression is associated with increased VEGF expression. **(A, B)** Western blot analysis of VEGFA protein expression in Timm44^OE^ MCF-7 cells, quantified as fold change over control (B), n = 3. (**C**) qRT-PCR analysis of VEGFA transcripts in Timm44^OE^ cells with or without MB-10 treatment, represented as fold change over controls. n = 3. **(D, E)** Immunoblot analysis of VEGFA protein levels in Timm44^OE^ cells following MB-10 treatment, further quantified as fold change over control (E), n = 3. **(F-H)** Western blot quantification of pan-AKT and ERK1/2 protein levels in control and Timm44^OE^ cells, further analyzed densitometrically (G, H), n = 3. (**I, J**) qRT-PCR–based analysis of ANGPT1 and ANGPT2 expression, represented as fold change over controls, n = 3. All data is presented as mean ± s.e.m. Two-tailed Student’s t-tests were used for pairwise comparisons between control and Timm44^OE^ cells, or Timm44^OE^ and MB-10–treated cells. *P < 0.05, **P < 0.01, ***P < 0.001, ****P < 0.0001.

To test whether this effect specifically stemmed from increased mitochondrial import capacity, we pharmacologically inhibited Timm44 activity using MB-10, a highly selective Timm44 inhibitor(*37*). Inhibition of Timm44 by MB-10 has been shown to disrupt TIM23-dependent protein translocation across the inner mitochondrial membrane (Miyata, Tang et al. 2017). Treatment of *Timm44^OE^* cells with MB-10 significantly reduced VEGFA levels (Fig. 1D, 1E), indicating a functional correlation between mitochondrial import capacity and heightened angiogenic output.

Because VEGFA frequently functions in an autocrine loop to enhance tumor cell proliferation by triggering downstream molecules such as ERK and AKT(*38, 39*), we examined the levels of Erk1/2 and pan-Akt which were found to be upregulated in *Timm44^OE^*cells (Fig. 1F-H). These observations also aligned with the pattern in cBioPortal datasets where Timm44-high tumors displayed strong correlation with MAP2K2 (MEK2) (Fig.S2D), principal upstream activators of the ERK. Consequent to activation of these VEGFA induced pro-growth factors, *Timm44^OE^* cells showed enhanced proliferation compared to controls (Fig. S2E). Beyond VEGFA, Timm44 overexpression also induced both ANGPT1 and ANGPT2 (Fig. 1I, J), that serve as critical modulators of endothelial activation and vessel stabilization. Interestingly, ANGPT1 and ANGPT2 displayed differential regulation. ANGPT1, which promotes vessel maturation(*40*), was modestly upregulated (Fig. 1J), whereas ANGPT2 known to facilitate sprouting and vascular permeability (*41, 42*) showed a more pronounced induction (Fig. 1I). This suggests that *Timm44^OE^* cells shift toward a pro-sprouting, pro-remodeling angiogenic profile typically associated with aggressive tumors. Together, these results indicate towards a potential link between mitochondrial translocase activity and endothelial growth signaling from tumor cells, positioning Timm44 as a central regulator.

### Timm44 activates endothelial sprouting and neovascularization

To determine whether the VEGFA and angiopoietins secreted by Timm44 overexpressing tumor cells were functionally capable of promoting neovascularization, we performed a 3D endothelial spheroid sprouting assay. Endothelial spheroids cultured in standard endothelial growth medium (EGM) displayed only minimal sprouts, reflecting their baseline angiogenic behavior. In contrast, conditioned media (CM) collected from *Timm44^OE^* breast cancer cells triggered a markedly more robust sprouting response than CM from control cells (Fig. 2A). Even when diluted to 75%, 50%, or 25% with EGM, the *Timm44^OE^* CM consistently triggered far more sprouting than control CM (Fig. 2A), indicating the presence of potent pro-angiogenic factors released by Timm44-high tumor cells. Quantitative measurements reinforced these observations since spheroids treated with *Timm44^OE^* CM displayed significantly more sprouts (Fig. 2B-E), longer average sprout lengths (Fig. 2F-I) at all dilutions. Additionally, the sprout diameter, a reflection of endothelial cord stability and lumen-forming potential, was substantially enhanced in spheroids exposed to higher CM concentrations (Fig. 2A *lower panel*). Notably, the average spheroid diameter itself was also significantly increased in spheroids treated with 50% and 100% CM derived from *Timm44^OE^* cells (Fig. 2J-K), indicating augmented endothelial growth and structural expansion under pro-angiogenic conditions. IL-6, used as a positive control for angiogenic stimulation, produced a similar response, supporting the strong angiogenic potency of Timm44-driven secreted factors. In parallel, *Timm44^OE^*CM also increased HUVEC proliferation overtime, relative to control CM at all tested dilutions (Fig 2L), reinforcing the idea that Timm44-dependent mitochondrial signaling drives a paracrine secretory program that supports multiple endothelial behaviors essential for angiogenesis.

**Fig 2.**
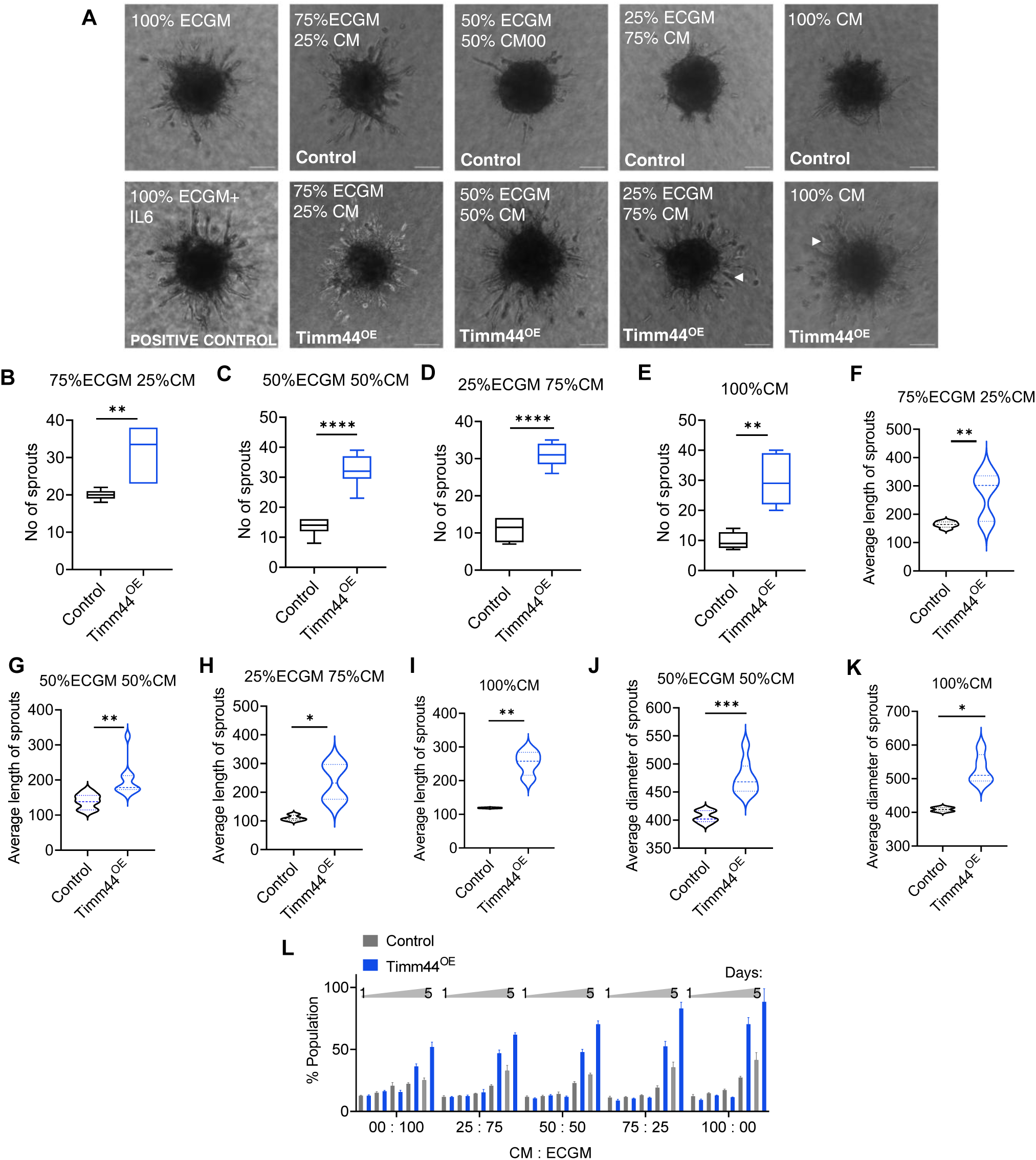
Timm44-high tumour cells promote endothelial growth and proliferation. **(A)** Endothelial spheroid sprouting assay performed using conditioned media (CM) from control and Timm44^OE^ cells at CM:ECGM ratios of 25:75, 50:50, 75:25, and 100:0, ECGM alone was used as the baseline control, while IL-6 + ECGM served as a positive control. **(B-E)** Quantitative analysis of endothelial sprout number. For each CM:ECGM ratios, the number of endothelial spheroids analysed (n) was as follows: 25:75 control (n = 7), Timm44^OE^ (n = 6); 50:50 control (n = 7), Timm44^OE^ (n = 10); 75:25 control (n = 4), Timm44^OE^ (n = 5); 100:0 control (n =4), Timm44^OE^ (n =5). **(F-I)** Quantitative analysis of average sprout length. The number of endothelial spheroids analysed (n) for each CM:ECGM ratio was as follows; (average sprout length represents the mean value per spheroid) 25:75 control (n = 6), Timm44^OE^ (n = 6); 50:50 control (n = 7), Timm44^OE^ (n = 10); 75:25 control (n = 4), Timm44^OE^ (n = 4); 100:0 control (n =2), Timm44^OE^ (n =4). **(J, K)** Quantitative analysis of endothelial sprout diameter. The number of endothelial spheroids analysed (n) was as follows; n = 5 (control) and 8 (Timm44^OE^) for 50:50, and n = 3 (control) and 5 (Timm44^OE^) for 100:0. Statistical significance of all quantifications, unless specifically mentioned, was assessed using two-tailed Student’s t-tests comparing control and Timm44^OE^ independently for each CM:ECGM ratio (25:75, 50:50, 75:25, and 100:0). **(L)** Endothelial cell proliferation assessed over a 5-day period following conditioned media treatment at the indicated ratios (n = 3). The sample with maximum growth rate was set at 100% and used for normalization of the rest (n=3). Statistical significance calculated using two-way Annova and P-value calculated for both row and column factor. *P < 0.05, **P < 0.01, ***P < 0.001, ****P < 0.0001.

To validate these effects in a more physiological setting, we used the chick chorioallantoic membrane (CAM) assay. Seventy-two hours after placing *Timm44^OE^* or control cells onto the CAM, embryos receiving Timm44-high cells showed a striking increase in vascular branching and density (Fig. S3A). The enhanced neovascularization observed in vivo closely mirrored the sprouting responses seen in vitro, reinforcing the idea that Timm44 actively promotes an angiogenic tumor phenotype. Together, these results form a functional bridge between our molecular observations and physiological angiogenesis.

### Timm44 mediated mitochondrial import activity orchestrates a coordinated biogenesis program

Since, Timm44 functions at the gateway of mitochondrial import machinery(*43*), we envisioned that its gain of function could result an increase in mitochondrial capacity by boosting its different components. We observed that *Timm44^OE^* mitochondria harboured high levels of TFAM (Fig. 3A, B) that is indispensable for mitochondrial DNA (mtDNA) replication, its packaging into nucleoids, and transcription initiation, acting as the key architectural factor that compacts mtDNA into nucleoids and recruits transcriptional polymerases(*44, 45*). Expectedly, the mtDNA copy number in *Timm44^OE^* mitochondria was ∼3-fold higher than the controls (Fig. 3C). Notably, treatment of *Timm44^OE^* cells with Timm44 inhibitor MB-10 reversed TFAM levels (Fig. S4A, 3B) and restored mtDNA copy number to those of control cells (Fig. 3C). This indicates that the increase in TFAM was a direct result of Timm44 activity at the import channel, which possibly facilitated the entry of nuclear-encoded factors that subsequently amplified mitochondrial transcription and replication. Therefore, we examined additional components of the mitochondrial gene-expression machinery and found significant upregulation of mitochondrial RNA polymerase (POLRMT)(*46*) (Fig. 3D), ATAD3 (Fig. 3E), a multifunctional regulator of nucleoid organization, cristae architecture, and mtDNA-protein interactions(*47–49*) and MRPL4 (Fig. 3F), a core large-subunit mitochondrial ribosomal protein essential for intra-mitochondrial translation; indicating an orchestrated enhancement of mitochondrial transcription and translation. Again, MB-10 treatment reversed these changes (Fig. 3D-F), strengthening the link between Timm44 activity and heightened mitochondrial transcriptional-translational output.

**Fig 3.**
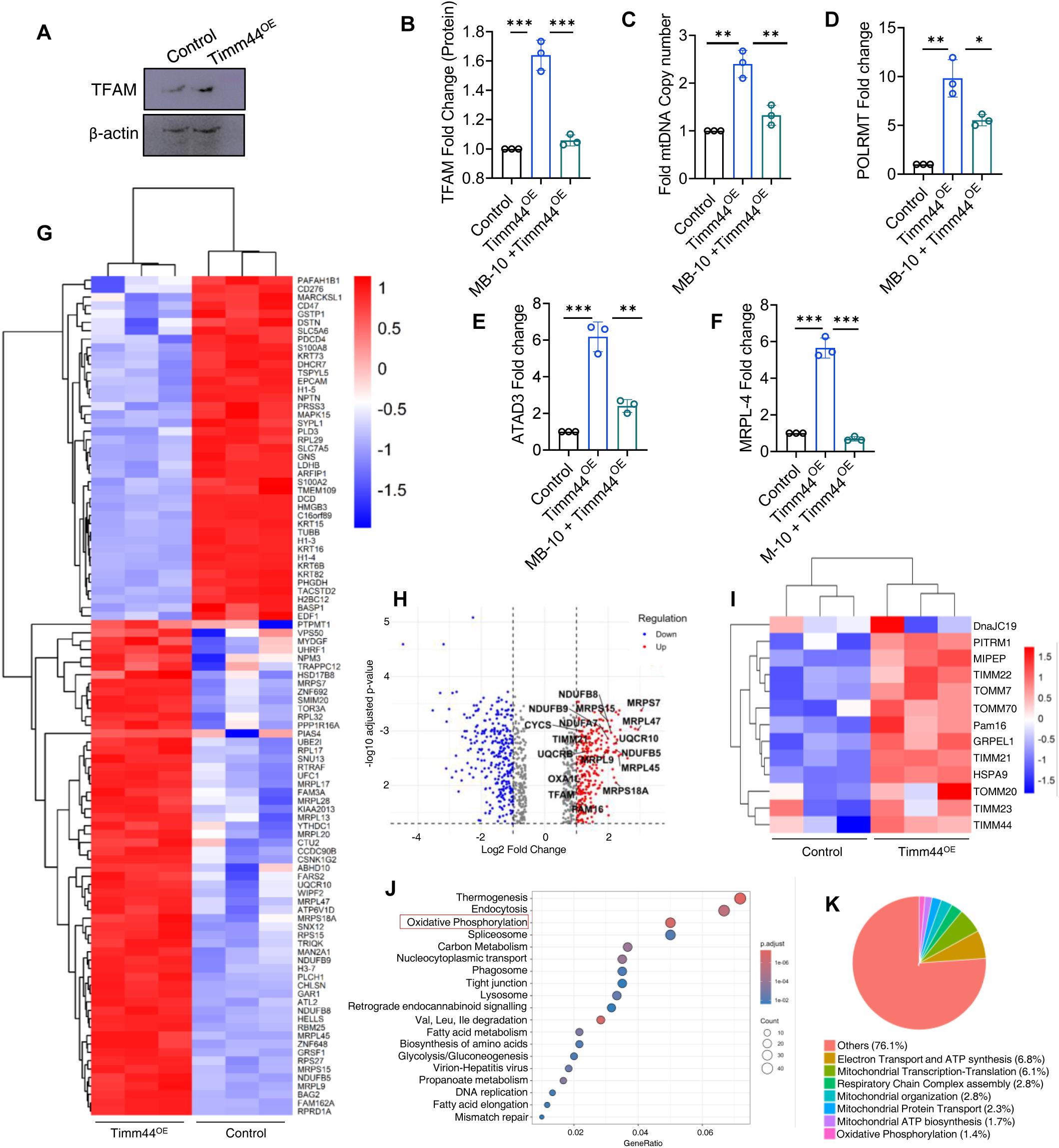
Protein-import based upregulation of mitochondria biogenesis. **(A, B)** Western blot analysis of TFAM protein levels in control and Timm44^OE^ cells, quantified, as fold change over controls. n = 3. Quantification of MB-10 treated sample was performed on S4A. **(C)** mtDNA copy number analysis calculated as fold change above control, for Timm44^OE^, and Timm44^OE^ MB-10 treated cells, n = 3. (**D-F**) qRT-PCR analysis of POLRMT, ATAD3, and MRPL4 mRNA levels in control, Timm44^OE^, and Timm44^OE^ MB-10 treated cells, represented as fold change over controls. n = 3. **(G)** Heatmap of the top 100 most differentially expressed genes selected based on variance. Gene expression values were log₂-transformed, Z-score normalized and displayed using a red-to-blue color scale as indicated. Both genes and samples were hierarchically clustered using Euclidean distance. **(H)** Volcano plot showing differential gene expression between the indicated conditions, with log₂ fold change on the x-axis and −log₁₀ (adjusted P value) on the y-axis. Significantly upregulated genes (adjusted P < 0.05, log₂ fold change > 1) and downregulated genes (adjusted P < 0.05, log₂ fold change < −1) are highlighted in red and blue, respectively. **(I)** Heatmap depicting the differential expression of mitochondrial import component genes. Expression values were Z-score normalized and shown using a red-to-blue color scale as indicated. Both genes and samples were hierarchically clustered using Euclidean distance. **(J)** Dot plot illustrating KEGG pathway enrichment analysis performed using the over-representation analysis (ORA) method for both upregulated and downregulated genes in Timm44^OE^ cells relative to control. Adjusted P values are represented by a color gradient ranging from red to blue, while the size of each dot corresponds to the number of genes enriched in each pathway. **(K)** Pie chart illustrating enriched upregulated biological processes in the proteomic dataset. For panel A-E, data is presented as mean ± s.e.m. Two-tailed Student’s t-tests were used for pairwise comparisons between control and Timm44^OE^ cells, or Timm44^OE^ and MB-10–treated cells. *P < 0.05, **P < 0.01, ***P < 0.001, ****P < 0.0001.

A more comprehensive view of mitochondrial remodeling was obtained through proteomic profiling of mitochondria-enriched fractions. In addition to elevated levels of inner mitochondrial membrane transport proteins such as Oxa1 and Timm21, which are known for assembly of respiratory subunits(*50, 51*), *Timm44^OE^* mitochondria showed a pronounced increase in proteins involved in mitochondrial ribosome assembly and gene expression (Fig. 3G, 3H). Consistent with these findings, volcano plot analysis (Fig. 3H) revealed significant enrichment of components of the mitochondrial translational machinery, along with mtDNA maintenance factors including TFAM, thereby, validating the above observations. We observed a coordinated upregulation of proteins associated with mitochondrial protein import, including multiple components of the TIM and TOM complexes, indicative of a broader reinforcement of the import machinery. This included prominent enrichment of direct Timm44 interactors, such as mortalin (HSPA9) and Pam16, underscoring co-regulation of the TIM23-motor complex (Fig. 3I). In parallel, a substantial fraction of proteins involved in oxidative phosphorylation was enriched in *Timm44^OE^* cells, pointing to a functional shift toward enhanced mitochondrial respiration (Fig. 3J, S4E). Approximately 24% of the upregulated proteins mapped to mitochondrial biogenesis and OXPHOS-related pathways, highlighting the coordinated nature of this mitochondrial activation program (Fig. 3K). Finally, to assess the clinical relevance of this signature, we analyzed TCGA BRCA datasets using cBioPortal. Timm44 expression displayed strong positive correlations with POLRMT, MRPL4, and ATAD3 (Fig. S4B-D), indicating that this transcriptional–biogenetic coupling observed experimentally is conserved across patient tumors. Together, these findings demonstrate that Timm44 gain-of-function reinforced the import machinery itself and initiated a coordinated mitochondrial biogenesis-like program encompassing mtDNA replication, transcription, translation, enrichment of mitochondrial ribosomal components and transcriptional regulators. This resulted into a functional remodeling of the organelle, raising the question whether this reprogramming culminated in a measurable enhancement of mitochondrial respiratory output and bioenergetic capacity.

### Timm44 couples mitochondrial protein import to enhanced Complex I-dependent respiratory capacity

To determine whether Timm44-driven mitochondrial remodeling translated into functional bioenergetic consequences, we next examined the mitochondrial respiratory performance. The *Timm44^OE^*cells showed minimal acidification of the extracellular media (Fig. S5A), overtime compared to controls and expectedly, exhibited much lower LDH expression (Fig. S5B), indicating a predominant reliance on mitochondrial respiration. We therefore, assessed the abundance of mitochondrial respiratory complex subunits and observed that *Timm44^OE^*cells showed significant upregulation of the nuclear-encoded components NDUFB8, UQCR2, and COXII, representing key subunits of Complexes I, III, and IV, respectively (Fig. 4A-C, S5C). In parallel, mitochondrial DNA-encoded components ND1 and CYTB, representing Complexes I and III respectively, showed increased transcript abundance (Fig. 4D, E), reflecting coordinated activation of respiratory chain components across nuclear and mitochondrial genomes, and aligning with our previous observations on enhanced nuclear-protein import and organeller transcription/translation output. Pharmacologic inhibition with MB-10 fully restored ND1 and CYTB expression to control levels (Fig. 4D, E), demonstrating that their induction is tightly coupled to Timm44-dependent import activity. Analysis of the mitochondrial proteome indicated cumulative upregulation of multiple subunits across all respiratory complexes in *Timm44^OE^* cells (Fig. 4F). To assess the physiological consequences of this molecular rewiring, we performed respirometry analysis (Fig. 4G). Timm44 overexpression broadly enhanced mitochondrial function, increasing basal (Fig. 4H) and maximal respiration (Fig. S5D), along with better spare respiratory capacity (Fig. 4I). The increase in mitochondrial respiration was mainly due to enhanced Complex I activity (Fig. S5E) and better Complex I-III transfer kinetics (Fig. S5F), along with parallel increase in Complex IV activity (Fig. S5G). The cells showed a shift towards on oxidative metabolism as indicated by very low non-mitochondrial respiration (Fig. 4J). Although the proton leak in *Timm44^OE^* mitochondria was high (Fig. 4K), probably due to high Complex I and Complex III activity, their coupling efficiency remained unchanged (Fig. 4L), indicating intact ETC performance. In fact, the cells showed higher RCR (Fig. 4M), indicating their better capability to couple oxygen consumption with ATP production (Fig. 4N). Hence, the *Timm44^OE^* mitochondria possessed better capability for ATP production and this functional enhancement correlated with higher intracellular ATP (Fig. 4O). Treatment with MB-10 normalized the ATP to levels equivalent to controls. The *Timm44^OE^* cells also showed a higher NAD⁺/NADH ratio (Fig. 4P), indicating utilization of these electron donors in a robust Complex I dependent respiration. Notably, mitochondrial mass (Fig. 4Q) and membrane potential (Fig. 4R) remained unchanged, and mitochondrial morphology (Fig. S5H) was preserved, suggesting that Timm44 enhances respiratory output without compromising structural integrity. PGC-1α expression (Fig. S5I, J) also remained unchanged in *Timm44^OE^* cells, indicating that the respiratory activation arises from enhanced import-driven mitochondrial transcription and translation rather than canonical biogenesis. Correlation analyses in TCGA BRCA datasets further showed strong associations between Timm44 and Complex I & III subunits (Fig. S5K, L). Collectively, these findings position Timm44 as a potent amplifier of respiratory capacity and mitochondrial energy production.

**Fig 4.**
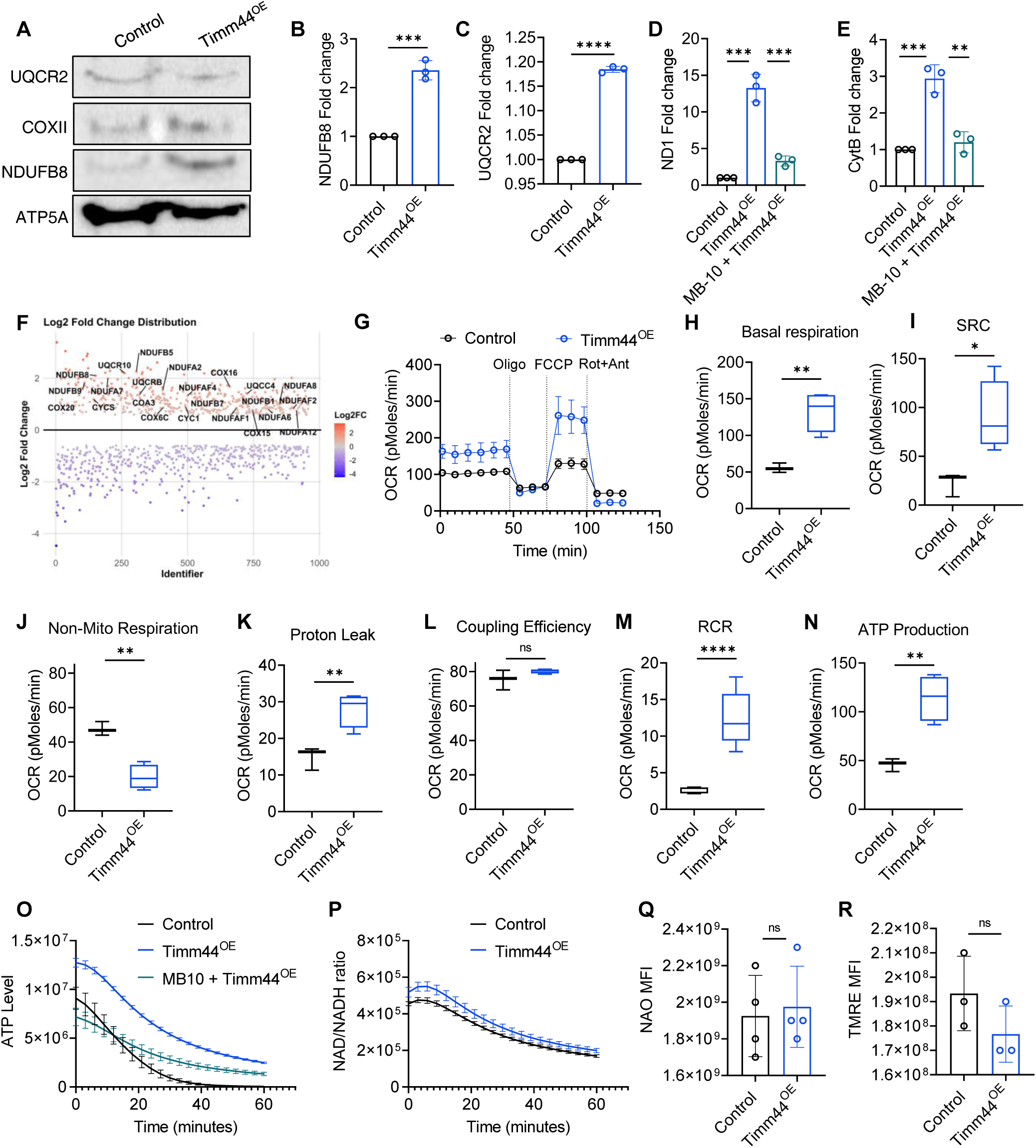
Enhancement of mitochondrial respiratory capacity by Timm44-induced biogenesis. **(A)** Immunoblot showing protein levels of representative nuclear-encoded respiratory chain subunits NDUFB8 (Complex I), COXII (Complex IV) and UQCR2 (Complex III) across control and Timm44^OE^ cells. **(B, C)** Densitometric quantification of NDUFB8 and UQCR2 protein levels normalized to loading control and represented as fold change over control, n=3. **(D, E)** qRT-PCR analysis of mitochondrial DNA-encoded respiratory subunits ND1 and CYTB across control, Timm44^OE^, and Timm44^OE^ MB-10–treated cells, calculated as fold change over controls, n = 3. **(F)** Scatter plot derived from mitochondrial proteomic analysis showing relative abundance of respiratory chain subunits across mitochondrial complexes. **(G-N)** Seahorse XF mitochondrial stress assay for control and Timm44^OE^ cells, normalized per μg of protein (Oligo- Oligomycin, Rot- Rotenone, Ant- Antimycin). The following respiratory parameters were calculated - basal and maximal respiration, spare respiratory capacity, non-mitochondrial respiration, coupling efficiency, respiratory control ratio (RCR), and ATP production, n = 3. **(O)** Intracellular ATP levels across control, Timm44^OE^, and Timm44^OE^ MB-10-treated cells, n=**(P)** NAD⁺/NADH ratio in control, Timm44^OE^, and Timm44^OE^ MB-10-treated cells, n = 3. **(Q, R)** NAO- and TMRE-based analysis of mitochondrial mass and membrane potential, n = All data are presented as mean ± SEM. Comparisons were performed between control and Timm44^OE^ cells, or between Timm44^OE^ and MB-10-treated Timm44^OE^ cells. Statistical significance was determined using two-tailed Student’s T-tests. *P < 0.05, **P < 0.01, ***P < 0.001, ****P < 0.0001.

### Mitochondrial hyperactivity via Timm44 drives Myc-centred proliferation

ETC hyperactivation due to enhanced activity of Complex I and III, resulted in an increase in superoxide levels in *Timm44^OE^* cells (Fig. 5A). This heightened oxidative state was associated with marked reduction in MnSOD (Fig. 5B, C), the primary mitochondrial enzyme responsible for dismutating O₂•⁻ into H₂O₂. GCL, the rate-limiting enzyme for glutathione synthesis, was also similarly downregulated (Fig. 5D, E), indicating diminished cellular capacity to detoxify mitochondrial ROS. In contrast, Prx3 (Fig. 5F, G) which detoxifies peroxides and GSK3β, a redox-responsive kinase linked to stress-induced apoptosis and mitophagy, remained largely unaltered(*52*) (Fig. 5H, 5I). To determine the functional consequences of this ROS-rich yet non-lethal environment, we assessed redox-sensitive regulators of proliferation. c-Myc, a canonical ROS-responsive transcriptional amplifier(*53, 54*), was strongly upregulated in *Timm44^OE^* cells, and its expression returned to baseline following MB-10 inhibition (Fig. 5J). Cdk1, a key Myc-cooperative regulator of cell-cycle progression(*55*), was similarly elevated and MB-10 sensitive (Fig. 5K). Beyond Myc and Cdk1, Timm44 overexpression induced a broader redox-responsive oncogenic factors comprising of PIN1 (Fig. 5L), a ROS-activated prolyl isomerase that stabilizes Myc(*56*); Cdc37 (Fig. 5M), an Hsp90 co-chaperone required for the maturation of multiple oncogenic kinases(*57*); and PPAN (Fig. 5N), a Myc-regulated factor essential for ribosome biogenesis and growth(*58*); and all of them normalized to controls levels upon MB-10 treatment. Consistent with activation of this mitogenic factor, *Timm44^OE^* cells proliferated faster than controls (Fig. S2E). cBioPortal analyses in TCGA-BRCA further revealed strong positive correlations between Timm44 and PIN1, Cdc37 and PPAN (Fig. S6A-C), demonstrating that this Timm44-linked redox signature is conserved across human tumors.

**Fig 5.**
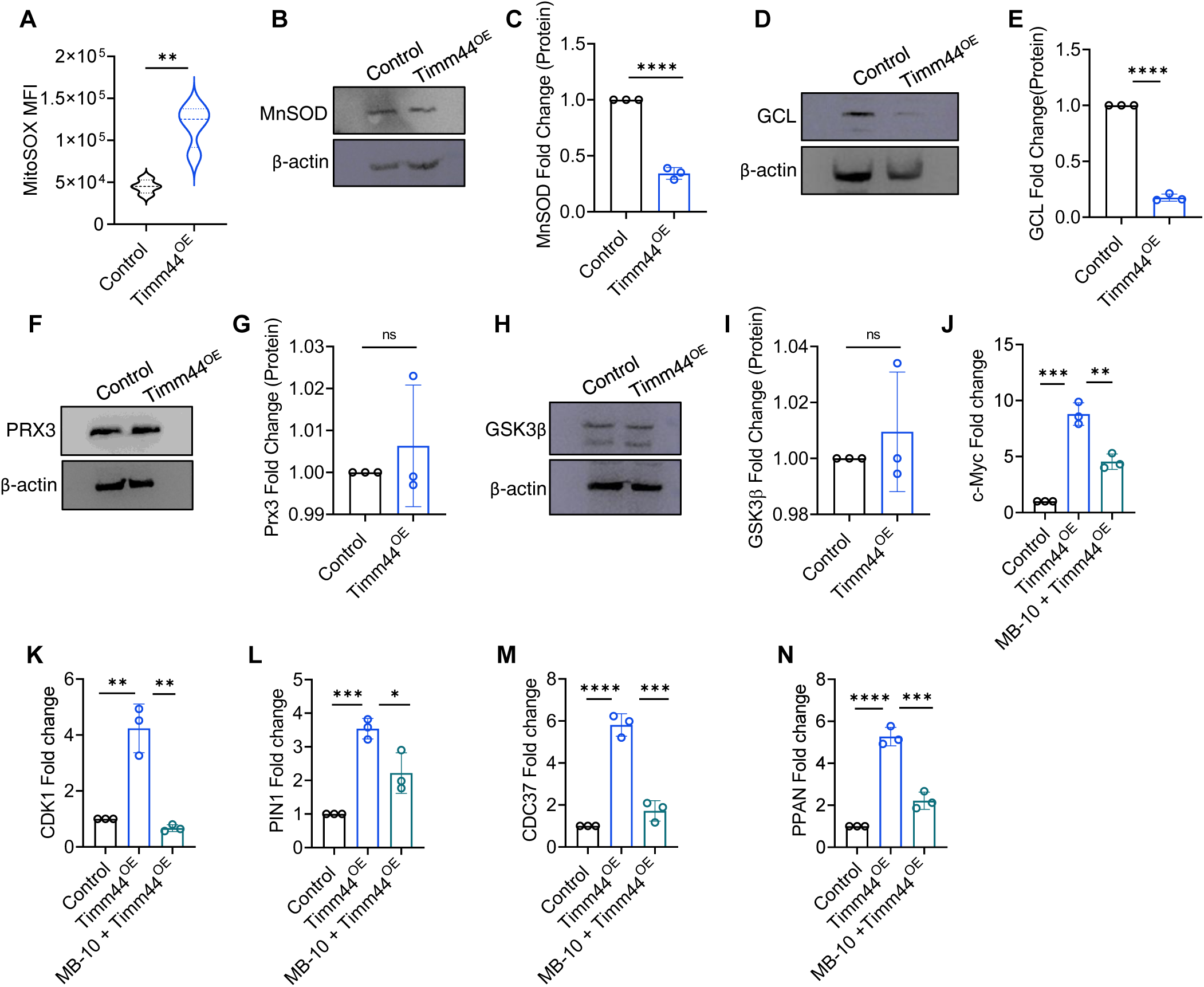
Alteration in redox-state by the respiration competent mitochondria. **(A)** Quantification of MitoSOX fluorescence intensity in control and Timm44^OE^ cells, n =4. **(B-I)** Immunoblot analysis and quantification of mitochondrial and cytosolic antioxidant components, including MnSOD (B, C), glutamate-cysteine ligase (GCL) (D, E), PRX3 (F, G) and GSK3β (H, I), with corresponding densitometric quantifications represented as fold-change over control, n= 3. **(J, K)** qRT–PCR analysis of c-MYC (J) and CDK1 (K) transcript levels calculated as fold change over control, n = 3. **(L-N)** qRT–PCR analysis of PIN1, CDC37, and PPAN expression in control, Timm44^OE^, and MB-10–treated cells, calculated as fold change over control, n= 3. All data are presented as mean ± SEM. Comparisons were performed between control and Timm44^OE^ cells, and Timm44^OE^ and MB-10–treated Timm44^OE^ cells. Statistical significance was determined using two-tailed Student’s t-tests. *P < 0.05, **P < 0.01, ***P < 0.001, ****P < 0.0001.

### Timm44-induced mitochondrial hyperfunction triggers HIF-independent VEGFA upregulation

Having established that Timm44 expressing cells show a robust dependence on aerobic respiration, we next sought to define how these metabolic changes culminate in robust VEGFA induction and the associated angiogenic phenotype. Typically, the vascularization is initiated by the master transcriptional factor, HIF-1α which is the canonical regulator of VEGFA under hypoxia and metabolic stress. We therefore, first asked whether Timm44 engages this pathway. Surprisingly, HIF-1α protein levels were significantly reduced in *Timm44^OE^* cells (Fig. 6B, S7A), indicating that VEGFA activation probably proceeds through a HIF-independent mechanism and necessitating identification of alternative transcriptional regulators. We systematically profiled transcription factors known to bind and activate the VEGFA promoter. We did not observe any change in the levels of STAT3(*59*) (Fig. 6C, S7B) and AP1 levels(*60*) (Fig. 6D). In contrast, SP1 transcript levels were markedly increased in *Timm44^OE^*cells and inhibition of Timm44 by MB-10 significantly reduced the levels of SP1 (Fig. 6E). To test whether elevated SP1 led to VEGFA induction, we depleted the SP1 transcript levels using RNAi which resulted in substantial reduction in VEGFA protein levels compared to *Timm44^OE^*(Fig. 6F, S7C) highlighting its role as an essential mediator of this transcriptional program.

**Fig 6.**
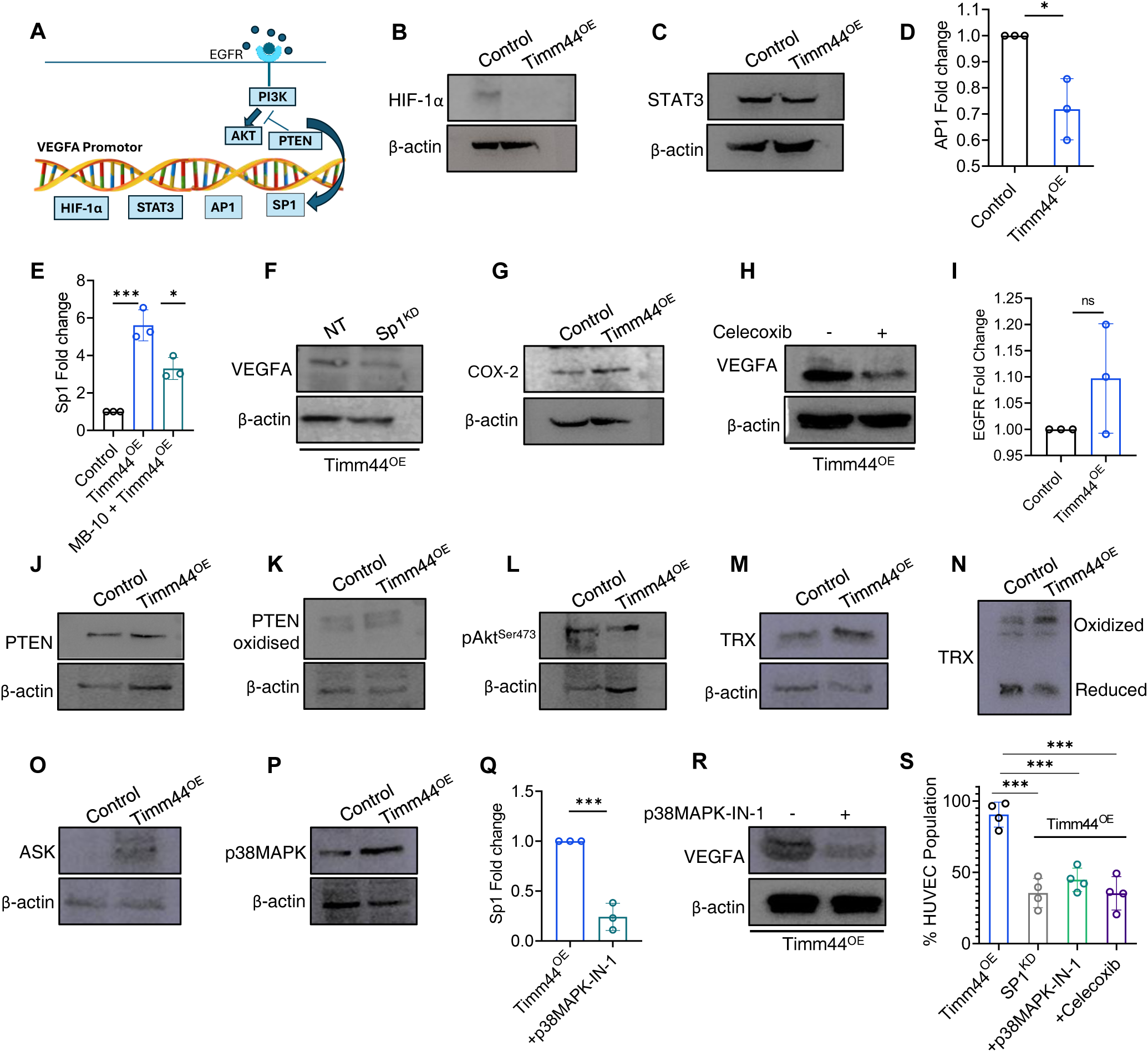
Mitochondrial ROS non-canonically induces VEGFA expression. **(A)** Schematic summarizing the major drivers of VEGFA expression. **(B)** Immunoblot analysis of HIF1α protein levels in control and Timm44^OE^ cells. **(C)** Immunoblot analysis of STAT3 protein levels. **(D, E)** qRT-PCR analysis of AP-1 (D) and SP1 (E) transcript levels calculated as fold change over control, n = 3. **(F)** Immunoblot analysis of VEGFA protein levels following shRNA-mediated SP1 depletion in Timm44^OE^ cells. **(G)** Immunoblot analysis of COX-2 protein expression. **(H)** Immunoblot analysis of VEGFA protein levels following pharmacological inhibition of COX-2 by Celecoxib. **(I)** qRT-PCR analysis of EGFR transcript abundance, calculated as fold change over control, n = 3. **(J, K)** Immunoblot analysis of total PTEN (J) and oxidized PTEN (K). **(L)** Levels of pAkt^Ser473^ estimated using immunoblotting. **(M, N)** Immunoblot analysis of thioredoxin (TRX) redox states under reducing (M) and non-reducing conditions (N) showing the oxidation states of the protein. **(O)** Immunoblot analysis of ASK1 protein levels. **(P)** Immunoblot analysis of p38 MAPK phosphorylation status. **(Q)** qRT-PCR analysis of SP1 transcript levels in *Timm44^OE^* cells treated with p38MAPK-IN-1 inhibitor, calculated as fold change over control, n = 3. **(R)** Immunoblot analysis of VEGFA protein levels following pharmacological inhibition of p38 MAPK by p38MAPK-IN-1 inhibitor. **(S)** Endothelial proliferation assay following SP1 silencing, COX-2 inhibition or p38 MAPK inhibition under Timm44-overexpressing condition. All data are presented as mean ± SEM. Comparisons were performed between control and Timm44^OE^ cells and Timm44^OE^ and MB-10–treated Timm44^OE^ cells or Timm44^OE^ cells and Timm44^OE^ cells transfected with SP1 siRNA/treated with an inhibitor (p38 MAPK-IN-1/celecoxib). Statistical significance was determined using two-tailed Student’s t-test. *P < 0.05, **P < 0.01, ***P < 0.001, ****P < 0.0001.

SP1 not only potentiates VEGFA transcription through direct promoter binding but is also known to engage the COX-2 promoter(*61*), thereby establishing a feed-forward regulatory loop that amplifies its own transcriptional output. Consistent with this idea, COX-2 expression was significantly elevated in *Timm44^OE^* cells (Fig. 6G, S7D). Pharmacological inhibition of COX-2 activity using Celecoxib resulted in a concomitant decrease in VEGFA expression (Fig. 6H, S7E), indicating that SP1 and COX-2 function cooperatively to sustain VEGFA induction.

EGFR-PI3K-Akt axis forms one of the predominant pathways that activate SP1. However, we did not find any change in EGFR levels(*62*) (Fig. 6I), which led us to probe whether PI3K(*63*) was activated by redox-inactivated PTEN. PTEN levels were found stable under reducing conditions (Fig. 6J, S7F), and non-reducing conditions did not indicate any change between the oxidised and reduced forms under Timm44^OE^ condition (Fig. 6K, S7G) thereby, ruling out the involvement of this axis. To further exclude a contribution of canonical PI3K-Akt-mTOR-dependent translational control to VEGFA upregulation(*64*), we examined the levels of pAkt^Ser473^ and phosphorylation status of 4EBP1, a key downstream effector of mTOR signaling. Notably, we found reduced accumulation of pAkt^Ser473^ in *Timm44^OE^* cells (Fig. 6L, S7H) and the levels of phosphorylated 4EBP1 in these cells remained unchanged compared to controls (Fig. S7I, J), indicating that enhanced VEGFA expression is unlikely to arise from increased cap-dependent translation. Given that *Timm44^OE^* cells exhibit elevated mitochondrial ROS as a consequence of ETC hyperactivation and disproportionate antioxidant buffering, and because p38MAPK is known to regulate SP1 stability and transcriptional activity(*65*), we reasoned that the p38MAPK might act as a connecting link between mitochondrial redox signaling to VEGFA induction in *Timm44^OE^*cells. Examination of TRX–ASK1–p38MAPK pathway(*66, 67*), indicated presence of high levels of TRX under reducing conditions (Fig. 6M, S7K), significant proportion of which was in the oxidized form as revealed by non-reducing gels (Fig. 6N, S7L); thus, indicating an oxidative shift that disrupts TRX-mediated inhibition of ASK1. Consistent with this relief of repression, ASK1 protein levels increased (Fig. 6O, S7M), and p38MAPK showed robust activation (Fig. 6P, S7N). Pharmacological inhibition of p38MAPK by p38MAPK-IN-1 led to a marked reduction in SP1 levels (Fig. 6Q) and VEGFA protein abundance (Fig. 6R, S7O), demonstrating that p38 activation is functionally necessary for Timm44-driven VEGFA upregulation. Finally, to assess whether the lowering of VEGFA levels translated to changes in endothelial growth phenotypes, we exposed the HUVEC cells to CM media derived from either SP1^RNAi^, Celecoxib or p38MAPK-IN-1 treated *Timm44^OE^* cells. Under all the above treatment conditions, the proliferative capacity of the endothelial cells was found to be weaker than the ones exposed to CM from untreated *Timm44^OE^* controls (Fig. 6S). Taken together, the results indicate how Timm44-promoted mitochondrial hyperactivation translated into a potent HIF-independent angiogenic transcriptional program, mediated by SP1 and triggered by redox-activated p38 MAPK.

## Discussion

Tumours sustain growth by inducing neovascularization or co-opting existing vessels(*68*). In healthy tissues, endothelial quiescence is maintained by a balance between pro- and anti-angiogenic signal(*12*). This equilibrium is disrupted during tumour progression by an angiogenic switch driven by oncogenic signalling(*69*), hypoxia-induced VEGF expression(*70*), and tumour-associated inflammation(*71*). Recent advances position mitochondria as central regulators of endothelial growth by integrating metabolic state, oxygen availability, and redox signalling(*72–75*). Under hypoxia, mitochondrial ROS stabilize HIF-1α and induce VEGF transcription(*76, 77*). Several mitochondrial regulators such as SIRT3, p66Shc, calcium transporters, fission protein DRP1, and Timm44 are essential for endothelial survival and inhibition of these collapses endothelial mitochondrial function and suppresses capillary sprouting(*78–83*). However, the role of tumour cell mitochondria in initiating neovascular growth is not known.

Highly proliferative tumours, stimulated by growth factors may not always coincide with hypoxic signalling. They exhibit metabolic reprogramming that favours oxidative phosphorylation (OXPHOS) and sustained biosynthesis to support rapid proliferation(*84–86*). Many tumours, particularly kidney cancer are dependent on Complex I-driven respiration for NAD+ regeneration and metastatic spread(*87*). Therefore, instead of passively responding to nutrient availability, cancer cells require activation of alternative strategies to ensure steady supply of precursors and essential metabolites through the vasculature. We therefore asked whether augmenting mitochondrial function in tumour cells activate an angiogenic program that is similar or distinct from the canonical HIF1α pathway.

Our results showed that tumour cells showing enhanced mitochondrial activity engaged an angiogenic program that was different from the canonical HIF-mediated process. We jumpstarted mitochondrial function by altering the import activity through TIM23-translocase. We overexpressed Timm44, the central scaffold protein that tethers the motor component of this translocase channel(*88*), reasoning that this would accelerate import of nuclear-encoded proteins into mitochondria and because this protein was highly expressed in many angiogenesis-dependent cancers. Indeed, Timm44 upregulation boosted mitochondrial gene expression machinery, as indicated by increased levels of mtDNA maintenance proteins and transcription/translation factors, leading to higher OXPHOS activity. In other words, cells ramped up oxidative capacity by improving import efficiency rather than by simply expanding organelle mass. This import-centric remodelling of mitochondria parallels recent insights that mitochondrial mRNAs are transcribed and turned over on vastly different timescales than nuclear ones(*89*), suggesting that altering import can rapidly modulate respiratory output.

The surge in respiratory flux due to high Complex I activity drove rapid reduction of ubiquinone to ubiquinol, feeding electrons into Complex III and generating semiquinone intermediates at the Qo site. These intermediates leak electrons to molecular oxygen generating superoxide(*90, 91*). Under our conditions this redox stress did not stabilize HIF-1α (no pseudohypoxia occurred), yet it strongly induced VEGFA expression and endothelial proliferation. Instead, mtROS oxidised the redox-sensitive Trx–Ask1 complex, releasing Ask1 from Trx inhibition. The freed active Ask1 phosphorylated p38MAPK which then promoted activation of transcription factor SP1. SP1, a zinc-finger factor(*92*) previously implicated in stress and inflammatory responses, drove transcription of VEGFA independently of HIF(*93*). Silencing SP1 severely impaired endothelial proliferation, consistent with its key role in angiogenic gene regulation.

In summary, we identify mitochondrial protein import-driven ETC hyperactivation as a previously unrecognized non-canonical determinant of tumour angiogenesis. By coupling respiratory activity with redox-sensitive kinases, and SP1-driven VEGF transcription, our findings indicate a hypoxia-independent, mitochondria-instructed angiogenic program. This work expands the angiogenesis paradigm beyond oxygen sensing, revealing that mitochondrial state itself can determine when and how tumours engage vascular support. Moving forward, it will be important to test the generality of this mechanism across tumour types and in physiologically relevant in vivo models, and to define how this pathway intersects with hypoxia- and inflammation-driven angiogenic signals. At the molecular level, future studies should address how rapidly mitochondrial import modulates nuclear OXPHOS gene expression and whether additional transcriptional regulators cooperate with SP1 in this redox-kinase context. From a translational perspective, our study may offer new strategies to disrupt angiogenesis, particularly in OXPHOS-dependent tumours.

## Acknowledgement

We heartily thank Ms. Rashi Gupta for her help in using the R-pipelines during plotting of proteomics data. We also thank NCCS, Pune for cell lines, Central Discovery Centre - BHU, DST-FIST and CAS facility of Department of Zoology. BHU for different instrumentation facilities.

## Author contributions

T.C., D.S conceived and designed experiments. T.C. performed majority of experiments and analysed data. S.M.B and T.C. performed spheroid assays under the supervision of M.J. P.P. and T.C. performed the proteomic studies. T.C. and D.S. wrote the manuscript.

## Data availability

The data that support the findings of this study are available within the main text and its Supplementary Information file. The lead contact will share all raw data associated with this paper upon reasonable request and, when applicable, fulfilment of appropriate material transfer agreements.

## References

1. P. Carmeliet, R. K. Jain, Angiogenesis in cancer and other diseases. Nature 407, 249–257 (2000).

2. D. Hanahan, R. A. Weinberg, Hallmarks of cancer: the next generation. Cell 144, 646–674 (2011).

3. J. Majidpoor, K. Mortezaee, Angiogenesis as a hallmark of solid tumors - clinical perspectives. Cell Oncol (Dordr) 44, 715–737 (2021).

4. N. Ferrara, Vascular endothelial growth factor and the regulation of angiogenesis. Recent Prog Horm Res 55, 15–35; discussion 35-16 (2000).

5. H. L. Goel, A. M. Mercurio, VEGF targets the tumour cell. Nat Rev Cancer 13, 871–882 (2013).

6. K. H. Plate, G. Breier, H. A. Weich, W. Risau, Vascular endothelial growth factor is a potential tumour angiogenesis factor in human gliomas in vivo. Nature 359, 845–848 (1992).

7. I. J. Fidler, The pathogenesis of cancer metastasis: the ‘seed and soil’ hypothesis revisited. Nat Rev Cancer 3, 453–458 (2003).

8. G. T. Motz, G. Coukos, Deciphering and reversing tumor immune suppression. Immunity 39, 61–73 (2013).

9. C. S. Abhinand, R. Raju, S. J. Soumya, P. S. Arya, P. R. Sudhakaran, VEGF-A/VEGFR2 signaling network in endothelial cells relevant to angiogenesis. J Cell Commun Signal 10, 347–354 (2016).

10. Y. Crawford, I. Kasman, L. Yu, C. Zhong, X. Wu, Z. Modrusan, J. Kaminker, N. Ferrara, PDGF-C mediates the angiogenic and tumorigenic properties of fibroblasts associated with tumors refractory to anti-VEGF treatment. Cancer Cell 15, 21–34 (2009).

11. M. Presta, P. Dell’Era, S. Mitola, E. Moroni, R. Ronca, M. Rusnati, Fibroblast growth factor/fibroblast growth factor receptor system in angiogenesis. Cytokine Growth Factor Rev 16, 159–178 (2005).

12. S. Kazerounian, J. Lawler, Integration of pro- and anti-angiogenic signals by endothelial cells. J Cell Commun Signal 12, 171–179 (2018).

13. P. H. Maxwell, M. S. Wiesener, G. W. Chang, S. C. Clifford, E. C. Vaux, M. E. Cockman, C. C. Wykoff, C. W. Pugh, E. R. Maher, P. J. Ratcliffe, The tumour suppressor protein VHL targets hypoxia-inducible factors for oxygen-dependent proteolysis. Nature 399, 271–275 (1999).

14. P. P. Kapitsinou, V. H. Haase, The VHL tumor suppressor and HIF: insights from genetic studies in mice. Cell Death Differ 15, 650–659 (2008).

15. H. Lu, R. A. Forbes, A. Verma, Hypoxia-inducible factor 1 activation by aerobic glycolysis implicates the Warburg effect in carcinogenesis. J Biol Chem 277, 23111–23115 (2002).

16. P. Sonveaux, F. Vegran, T. Schroeder, M. C. Wergin, J. Verrax, Z. N. Rabbani, C. J. De Saedeleer, K. M. Kennedy, C. Diepart, B. F. Jordan, M. J. Kelley, B. Gallez, M. L. Wahl, O. Feron, M. W. Dewhirst, Targeting lactate-fueled respiration selectively kills hypoxic tumor cells in mice. J Clin Invest 118, 3930–3942 (2008).

17. W. G. Kaelin, Jr., P. J. Ratcliffe, Oxygen sensing by metazoans: the central role of the HIF hydroxylase pathway. Mol Cell 30, 393–402 (2008).

18. R. J. DeBerardinis, A. Mancuso, E. Daikhin, I. Nissim, M. Yudkoff, S. Wehrli, C. B. Thompson, Beyond aerobic glycolysis: transformed cells can engage in glutamine metabolism that exceeds the requirement for protein and nucleotide synthesis. Proc Natl Acad Sci U S A 104, 19345–19350 (2007).

19. L. B. Sullivan, D. Y. Gui, M. G. Vander Heiden, Altered metabolite levels in cancer: implications for tumour biology and cancer therapy. Nat Rev Cancer 16, 680–693 (2016).

20. V. S. LeBleu, J. T. O’Connell, K. N. Gonzalez Herrera, H. Wikman, K. Pantel, M. C. Haigis, F. M. de Carvalho, A. Damascena, L. T. Domingos Chinen, R. M. Rocha, J. M. Asara, R. Kalluri, PGC-1alpha mediates mitochondrial biogenesis and oxidative phosphorylation in cancer cells to promote metastasis. Nat Cell Biol 16, 992–1003, 1001-1015 (2014).

21. I. Martinez-Reyes, N. S. Chandel, Mitochondrial TCA cycle metabolites control physiology and disease. Nat Commun 11, 102 (2020).

22. K. De Bock, M. Georgiadou, P. Carmeliet, Role of endothelial cell metabolism in vessel sprouting. Cell Metab 18, 634–647 (2013).

23. K. De Bock, M. Georgiadou, S. Schoors, A. Kuchnio, B. W. Wong, A. R. Cantelmo, A. Quaegebeur, B. Ghesquiere, S. Cauwenberghs, G. Eelen, L. K. Phng, I. Betz, B. Tembuyser, K. Brepoels, J. Welti, I. Geudens, I. Segura, B. Cruys, F. Bifari, I. Decimo, R. Blanco, S. Wyns, J. Vangindertael, S. Rocha, R. T. Collins, S. Munck, D. Daelemans, H. Imamura, R. Devlieger, M. Rider, P. P. Van Veldhoven, F. Schuit, R. Bartrons, J. Hofkens, P. Fraisl, S. Telang, R. J. Deberardinis, L. Schoonjans, S. Vinckier, J. Chesney, H. Gerhardt, M. Dewerchin, P. Carmeliet, Role of PFKFB3-driven glycolysis in vessel sprouting. Cell 154, 651–663 (2013).

24. H. Huang, S. Vandekeere, J. Kalucka, L. Bierhansl, A. Zecchin, U. Bruning, A. Visnagri, N. Yuldasheva, J. Goveia, B. Cruys, K. Brepoels, S. Wyns, S. Rayport, B. Ghesquiere, S. Vinckier, L. Schoonjans, R. Cubbon, M. Dewerchin, G. Eelen, P. Carmeliet, Role of glutamine and interlinked asparagine metabolism in vessel formation. EMBO J 36, 2334–2352 (2017).

25. S. Schoors, U. Bruning, R. Missiaen, K. C. Queiroz, G. Borgers, I. Elia, A. Zecchin, A. R. Cantelmo, S. Christen, J. Goveia, W. Heggermont, L. Godde, S. Vinckier, P. P. Van Veldhoven, G. Eelen, L. Schoonjans, H. Gerhardt, M. Dewerchin, M. Baes, K. De Bock, B. Ghesquiere, S. Y. Lunt, S. M. Fendt, P. Carmeliet, Fatty acid carbon is essential for dNTP synthesis in endothelial cells. Nature 520, 192–197 (2015).

26. D. Mokranjac, W. Neupert, The many faces of the mitochondrial TIM23 complex. Biochim Biophys Acta 1797, 1045–1054 (2010).

27. S. Y. Ting, B. A. Schilke, M. Hayashi, E. A. Craig, Architecture of the TIM23 inner mitochondrial translocon and interactions with the matrix import motor. J Biol Chem 289, 28689–28696 (2014).

28. N. Jain, A. Chacinska, P. Rehling, Understanding mitochondrial protein import: a revised model of the presequence translocase. Trends Biochem Sci 50, 585–595 (2025).

29. T. Krimmer, J. Rassow, W. H. Kunau, W. Voos, N. Pfanner, Mitochondrial protein import motor: the ATPase domain of matrix Hsp70 is crucial for binding to Tim44, while the peptide binding domain and the carboxy-terminal segment play a stimulatory role. Mol Cell Biol 20, 5879–5887 (2000).

30. F. Moro, K. Okamoto, M. Donzeau, W. Neupert, M. Brunner, Mitochondrial protein import: molecular basis of the ATP-dependent interaction of MtHsp70 with Tim44. J Biol Chem 277, 6874–6880 (2002).

31. Z. R. Ma, H. P. Li, S. Z. Cai, S. Y. Du, X. Chen, J. Yao, X. Cao, Y. F. Zhen, Q. Wang, The mitochondrial protein TIMM44 is required for angiogenesis in vitro and in vivo. Cell Death Dis 14, 307 (2023).

32. A. Merlin, W. Voos, A. C. Maarse, M. Meijer, N. Pfanner, J. Rassow, The J-related segment of tim44 is essential for cell viability: a mutant Tim44 remains in the mitochondrial import site, but inefficiently recruits mtHsp70 and impairs protein translocation. J Cell Biol 145, 961–972 (1999).

33. Y. Wang, A. Katayama, T. Terami, X. Han, T. Nunoue, D. Zhang, S. Teshigawara, J. Eguchi, A. Nakatsuka, K. Murakami, D. Ogawa, Y. Furuta, H. Makino, J. Wada, Translocase of inner mitochondrial membrane 44 alters the mitochondrial fusion and fission dynamics and protects from type 2 diabetes. Metabolism 64, 677–688 (2015).

34. Y. Zhang, J. Wada, I. Hashimoto, J. Eguchi, A. Yasuhara, Y. S. Kanwar, K. Shikata, H. Makino, Therapeutic approach for diabetic nephropathy using gene delivery of translocase of inner mitochondrial membrane 44 by reducing mitochondrial superoxide production. J Am Soc Nephrol 17, 1090–1101 (2006).

35. D. C. Wallace, Mitochondria and cancer. Nat Rev Cancer 12, 685–698 (2012).

36. D. Schiller, Y. C. Cheng, Q. Liu, W. Walter, E. A. Craig, Residues of Tim44 involved in both association with the translocon of the inner mitochondrial membrane and regulation of mitochondrial Hsp70 tethering. Mol Cell Biol 28, 4424–4433 (2008).

37. N. Miyata, Z. Tang, M. A. Conti, M. E. Johnson, C. J. Douglas, S. A. Hasson, R. Damoiseaux, C. A. Chang, C. M. Koehler, Adaptation of a Genetic Screen Reveals an Inhibitor for Mitochondrial Protein Import Component Tim44. J Biol Chem 292, 5429–5442 (2017).

38. Y. Lin, E. Zhai, B. Liao, L. Xu, X. Zhang, S. Peng, Y. He, S. Cai, Z. Zeng, M. Chen, Autocrine VEGF signaling promotes cell proliferation through a PLC-dependent pathway and modulates Apatinib treatment efficacy in gastric cancer. Oncotarget 8, 11990–12002 (2017).

39. M. Simons, E. Gordon, L. Claesson-Welsh, Mechanisms and regulation of endothelial VEGF receptor signalling. Nat Rev Mol Cell Biol 17, 611–625 (2016).

40. S. Davis, T. H. Aldrich, P. F. Jones, A. Acheson, D. L. Compton, V. Jain, T. E. Ryan, J. Bruno, C. Radziejewski, P. C. Maisonpierre, G. D. Yancopoulos, Isolation of angiopoietin-1, a ligand for the TIE2 receptor, by secretion-trap expression cloning. Cell 87, 1161–1169 (1996).

41. T. Asahara, D. Chen, T. Takahashi, K. Fujikawa, M. Kearney, M. Magner, G. D. Yancopoulos, J. M. Isner, Tie2 receptor ligands, angiopoietin-1 and angiopoietin-2, modulate VEGF-induced postnatal neovascularization. Circ Res 83, 233–240 (1998).

42. H. Huang, A. Bhat, G. Woodnutt, R. Lappe, Targeting the ANGPT-TIE2 pathway in malignancy. Nat Rev Cancer 10, 575–585 (2010).

43. D. P. Hutu, B. Guiard, A. Chacinska, D. Becker, N. Pfanner, P. Rehling, M. van der Laan, Mitochondrial protein import motor: differential role of Tim44 in the recruitment of Pam17 and J-complex to the presequence translocase. Mol Biol Cell 19, 2642–2649 (2008).

44. M. I. Ekstrand, M. Falkenberg, A. Rantanen, C. B. Park, M. Gaspari, K. Hultenby, P. Rustin, C. M. Gustafsson, N. G. Larsson, Mitochondrial transcription factor A regulates mtDNA copy number in mammals. Hum Mol Genet 13, 935–944 (2004).

45. N. G. Larsson, J. Wang, H. Wilhelmsson, A. Oldfors, P. Rustin, M. Lewandoski, G. S. Barsh, D. A. Clayton, Mitochondrial transcription factor A is necessary for mtDNA maintenance and embryogenesis in mice. Nat Genet 18, 231–236 (1998).

46. J. J. Arnold, E. D. Smidansky, I. M. Moustafa, C. E. Cameron, Human mitochondrial RNA polymerase: structure-function, mechanism and inhibition. Biochim Biophys Acta 1819, 948–960 (2012).

47. J. M. Gerhold, S. Cansiz-Arda, M. Lohmus, O. Engberg, A. Reyes, H. van Rennes, A. Sanz, I. J. Holt, H. M. Cooper, J. N. Spelbrink, Human Mitochondrial DNA-Protein Complexes Attach to a Cholesterol-Rich Membrane Structure. Sci Rep 5, 15292 (2015).

48. J. He, H. M. Cooper, A. Reyes, M. Di Re, H. Sembongi, T. R. Litwin, J. Gao, K. C. Neuman, I. M. Fearnley, A. Spinazzola, J. E. Walker, I. J. Holt, Mitochondrial nucleoid interacting proteins support mitochondrial protein synthesis. Nucleic Acids Res 40, 6109–6121 (2012).

49. C. Kukat, N. G. Larsson, mtDNA makes a U-turn for the mitochondrial nucleoid. Trends Cell Biol 23, 457–463 (2013).

50. N. Altamura, N. Capitanio, N. Bonnefoy, S. Papa, G. Dujardin, The Saccharomyces cerevisiae OXA1 gene is required for the correct assembly of cytochrome c oxidase and oligomycin-sensitive ATP synthase. FEBS Lett 382, 111–115 (1996).

51. D. U. Mick, S. Dennerlein, H. Wiese, R. Reinhold, D. Pacheu-Grau, I. Lorenzi, F. Sasarman, W. Weraarpachai, E. A. Shoubridge, B. Warscheid, P. Rehling, MITRAC links mitochondrial protein translocation to respiratory-chain assembly and translational regulation. Cell 151, 1528–1541 (2012).

52. P. Watcharasit, A. Thiantanawat, J. Satayavivad, GSK3 promotes arsenite-induced apoptosis via facilitation of mitochondria disruption. J Appl Toxicol 28, 466–474 (2008).

53. B. Benassi, M. Fanciulli, F. Fiorentino, A. Porrello, G. Chiorino, M. Loda, G. Zupi, A. Biroccio, c-Myc phosphorylation is required for cellular response to oxidative stress. Mol Cell 21, 509–519 (2006).

54. C. Y. Lin, J. Loven, P. B. Rahl, R. M. Paranal, C. B. Burge, J. E. Bradner, T. I. Lee, R. A. Young, Transcriptional amplification in tumor cells with elevated c-Myc. Cell 151, 56–67 (2012).

55. Y. Yang, K. Xue, Z. Li, W. Zheng, W. Dong, J. Song, S. Sun, T. Ma, W. Li, [Corrigendum] c-Myc regulates the CDK1/cyclin B1 dependent-G2/M cell cycle progression by histone H4 acetylation in Raji cells. Int J Mol Med 44, 1988 (2019).

56. A. S. Farrell, C. Pelz, X. Wang, C. J. Daniel, Z. Wang, Y. Su, M. Janghorban, X. Zhang, C. Morgan, S. Impey, R. C. Sears, Pin1 regulates the dynamics of c-Myc DNA binding to facilitate target gene regulation and oncogenesis. Mol Cell Biol 33, 2930–2949 (2013).

57. J. R. Smith, P. A. Clarke, E. de Billy, P. Workman, Silencing the cochaperone CDC37 destabilizes kinase clients and sensitizes cancer cells to HSP90 inhibitors. Oncogene 28, 157–169 (2009).

58. N. Zielke, A. Vaharautio, J. Liu, T. Kivioja, J. Taipale, Upregulation of ribosome biogenesis via canonical E-boxes is required for Myc-driven proliferation. Dev Cell 57, 1024–1036 e1025 (2022).

59. G. Niu, K. L. Wright, M. Huang, L. Song, E. Haura, J. Turkson, S. Zhang, T. Wang, D. Sinibaldi, D. Coppola, R. Heller, L. M. Ellis, J. Karras, J. Bromberg, D. Pardoll, R. Jove, H. Yu, Constitutive Stat3 activity up-regulates VEGF expression and tumor angiogenesis. Oncogene 21, 2000–2008 (2002).

60. J. Jia, T. Ye, P. Cui, Q. Hua, H. Zeng, D. Zhao, AP-1 transcription factor mediates VEGF-induced endothelial cell migration and proliferation. Microvasc Res 105, 103–108 (2016).

61. H. Hu, T. Han, M. Zhuo, L. L. Wu, C. Yuan, L. Wu, W. Lei, F. Jiao, L. W. Wang, Elevated COX-2 Expression Promotes Angiogenesis Through EGFR/p38-MAPK/Sp1-Dependent Signalling in Pancreatic Cancer. Sci Rep 7, 470 (2017).

62. N. Pore, Z. Jiang, A. Gupta, G. Cerniglia, G. D. Kao, A. Maity, EGFR tyrosine kinase inhibitors decrease VEGF expression by both hypoxia-inducible factor (HIF)-1-independent and HIF-1-dependent mechanisms. Cancer Res 66, 3197–3204 (2006).

63. S. B. Choi, J. B. Park, T. J. Song, S. Y. Choi, Molecular mechanism of HIF-1-independent VEGF expression in a hepatocellular carcinoma cell line. Int J Mol Med 28, 449–454 (2011).

64. S. Braunstein, K. Karpisheva, C. Pola, J. Goldberg, T. Hochman, H. Yee, J. Cangiarella, R. Arju, S. C. Formenti, R. J. Schneider, A hypoxia-controlled cap-dependent to cap-independent translation switch in breast cancer. Mol Cell 28, 501–512 (2007).

65. B. A. Rose, T. Yokota, V. Chintalgattu, S. Ren, L. Iruela-Arispe, A. Y. Khakoo, S. Minamisawa, Y. Wang, Cardiac myocyte p38alpha kinase regulates angiogenesis via myocyte-endothelial cell cross-talk during stress-induced remodeling in the heart. J Biol Chem 292, 12787–12800 (2017).

66. M. Saitoh, H. Nishitoh, M. Fujii, K. Takeda, K. Tobiume, Y. Sawada, M. Kawabata, K. Miyazono, H. Ichijo, Mammalian thioredoxin is a direct inhibitor of apoptosis signal-regulating kinase (ASK) 1. EMBO J 17, 2596–2606 (1998).

67. K. Tobiume, A. Matsuzawa, T. Takahashi, H. Nishitoh, K. Morita, K. Takeda, O. Minowa, K. Miyazono, T. Noda, H. Ichijo, ASK1 is required for sustained activations of JNK/p38 MAP kinases and apoptosis. EMBO Rep 2, 222–228 (2001).

68. J. Holash, P. C. Maisonpierre, D. Compton, P. Boland, C. R. Alexander, D. Zagzag, G. D. Yancopoulos, S. J. Wiegand, Vessel cooption, regression, and growth in tumors mediated by angiopoietins and VEGF. Science 284, 1994–1998 (1999).

69. D. Hanahan, J. Folkman, Patterns and emerging mechanisms of the angiogenic switch during tumorigenesis. Cell 86, 353–364 (1996).

70. D. Shweiki, M. Neeman, A. Itin, E. Keshet, Induction of vascular endothelial growth factor expression by hypoxia and by glucose deficiency in multicell spheroids: implications for tumor angiogenesis. Proc Natl Acad Sci U S A 92, 768–772 (1995).

71. L. M. Coussens, Z. Werb, Inflammation and cancer. Nature 420, 860–867 (2002).

72. S. Caja, J. A. Enriquez, Mitochondria in endothelial cells: Sensors and integrators of environmental cues. Redox Biol 12, 821–827 (2017).

73. L. P. Diebold, H. J. Gil, P. Gao, C. A. Martinez, S. E. Weinberg, N. S. Chandel, Mitochondrial complex III is necessary for endothelial cell proliferation during angiogenesis. Nat Metab 1, 158–171 (2019).

74. D. Park, P. J. Dilda, Mitochondria as targets in angiogenesis inhibition. Mol Aspects Med 31, 113–131 (2010).

75. L. M. Schiffmann, J. P. Werthenbach, F. Heintges-Kleinhofer, J. M. Seeger, M. Fritsch, S. D. Gunther, S. Willenborg, S. Brodesser, C. Lucas, C. Jungst, M. C. Albert, F. Schorn, A. Witt, C. T. Moraes, C. J. Bruns, M. Pasparakis, M. Kronke, S. A. Eming, O. Coutelle, H. Kashkar, Mitochondrial respiration controls neoangiogenesis during wound healing and tumour growth. Nat Commun 11, 3653 (2020).

76. N. S. Chandel, E. Maltepe, E. Goldwasser, C. E. Mathieu, M. C. Simon, P. T. Schumacker, Mitochondrial reactive oxygen species trigger hypoxia-induced transcription. Proc Natl Acad Sci U S A 95, 11715–11720 (1998).

77. A. Reichard, K. Asosingh, The role of mitochondria in angiogenesis. Mol Biol Rep 46, 1393–1400 (2019).

78. B. R. Alevriadou, S. Shanmughapriya, A. Patel, P. B. Stathopulos, M. Madesh, Mitochondrial Ca(2+) transport in the endothelium: regulation by ions, redox signalling and mechanical forces. J R Soc Interface 14, (2017).

79. A. Csiszar, N. Labinskyy, J. T. Pinto, P. Ballabh, H. Zhang, G. Losonczy, K. Pearson, R. de Cabo, P. Pacher, C. Zhang, Z. Ungvari, Resveratrol induces mitochondrial biogenesis in endothelial cells. Am J Physiol Heart Circ Physiol 297, H13–20 (2009).

80. J. Y. Jin, X. X. Wei, X. L. Zhi, X. H. Wang, D. Meng, Drp1-dependent mitochondrial fission in cardiovascular disease. Acta Pharmacol Sin 42, 655–664 (2021).

81. S. Kumar, P66Shc and vascular endothelial function. Biosci Rep 39, (2019).

82. S. Liu, Y. Zhao, H. Yao, L. Zhang, C. Chen, Z. Zheng, S. Jin, DRP1 knockdown and atorvastatin alleviate ox-LDL-induced vascular endothelial cells injury: DRP1 is a potential target for preventing atherosclerosis. Exp Cell Res 429, 113688 (2023).

83. J. Oshikawa, S. J. Kim, E. Furuta, C. Caliceti, G. F. Chen, R. D. McKinney, F. Kuhr, I. Levitan, T. Fukai, M. Ushio-Fukai, Novel role of p66Shc in ROS-dependent VEGF signaling and angiogenesis in endothelial cells. Am J Physiol Heart Circ Physiol 302, H724–732 (2012).

84. L. Ippolito, A. Marini, L. Cavallini, A. Morandi, L. Pietrovito, G. Pintus, E. Giannoni, T. Schrader, M. Puhr, P. Chiarugi, M. L. Taddei, Metabolic shift toward oxidative phosphorylation in docetaxel resistant prostate cancer cells. Oncotarget 7, 61890–61904 (2016).

85. S. Rodriguez-Enriquez, P. A. Vital-Gonzalez, F. L. Flores-Rodriguez, A. Marin-Hernandez, L. Ruiz-Azuara, R. Moreno-Sanchez, Control of cellular proliferation by modulation of oxidative phosphorylation in human and rodent fast-growing tumor cells. Toxicol Appl Pharmacol 215, 208–217 (2006).

86. A. S. Tan, J. W. Baty, M. V. Berridge, The role of mitochondrial electron transport in tumorigenesis and metastasis. Biochim Biophys Acta 1840, 1454–1463 (2014).

87. D. Bezwada, L. Perelli, N. P. Lesner, L. Cai, B. Brooks, Z. Wu, H. S. Vu, V. Sondhi, D. L. Cassidy, S. Kasitinon, S. Kelekar, F. Cai, A. B. Aurora, M. Patrick, A. Leach, R. Ghandour, Y. Zhang, D. Do, P. McDaniel, J. Sudderth, D. Dumesnil, S. House, T. Rosales, A. M. Poole, Y. Lotan, S. Woldu, A. Bagrodia, X. Meng, J. A. Cadeddu, P. Mishra, J. Garcia-Bermudez, I. Pedrosa, P. Kapur, K. D. Courtney, C. R. Malloy, G. Genovese, V. Margulis, R. J. DeBerardinis, Mitochondrial complex I promotes kidney cancer metastasis. Nature 633, 923–931 (2024).

88. U. Bomer, A. C. Maarse, F. Martin, A. Geissler, A. Merlin, B. Schonfisch, M. Meijer, N. Pfanner, J. Rassow, Separation of structural and dynamic functions of the mitochondrial translocase: Tim44 is crucial for the inner membrane import sites in translocation of tightly folded domains, but not of loosely folded preproteins. EMBO J 17, 4226–4237 (1998).

89. E. McShane, M. Couvillion, R. Ietswaart, G. Prakash, B. M. Smalec, I. Soto, A. R. Baxter-Koenigs, K. Choquet, L. S. Churchman, A kinetic dichotomy between mitochondrial and nuclear gene expression processes. Mol Cell 84, 1541–1555 e1511 (2024).

90. F. L. Muller, Y. Liu, H. Van Remmen, Complex III releases superoxide to both sides of the inner mitochondrial membrane. J Biol Chem 279, 49064–49073 (2004).

91. C. L. Quinlan, I. V. Perevoshchikova, M. Hey-Mogensen, A. L. Orr, M. D. Brand, Sites of reactive oxygen species generation by mitochondria oxidizing different substrates. Redox Biol 1, 304–312 (2013).

92. J. Kaczynski, T. Cook, R. Urrutia, Sp1- and Kruppel-like transcription factors. Genome Biol 4, 206 (2003).

93. N. Pore, S. Liu, H. K. Shu, B. Li, D. Haas-Kogan, D. Stokoe, J. Milanini-Mongiat, G. Pages, D. M. O’Rourke, E. Bernhard, A. Maity, Sp1 is involved in Akt-mediated induction of VEGF expression through an HIF-1-independent mechanism. Mol Biol Cell 15, 4841–4853 (2004).

