## Supplementary Figures for "Protein import-driven mitochondrial hyperactivation dictates angiogenic signalling independently of HIF-1α"

**Figure S1**

A

Expression levels of Timm44 across tumor and normal tissues in TCGA and GTEx datasets

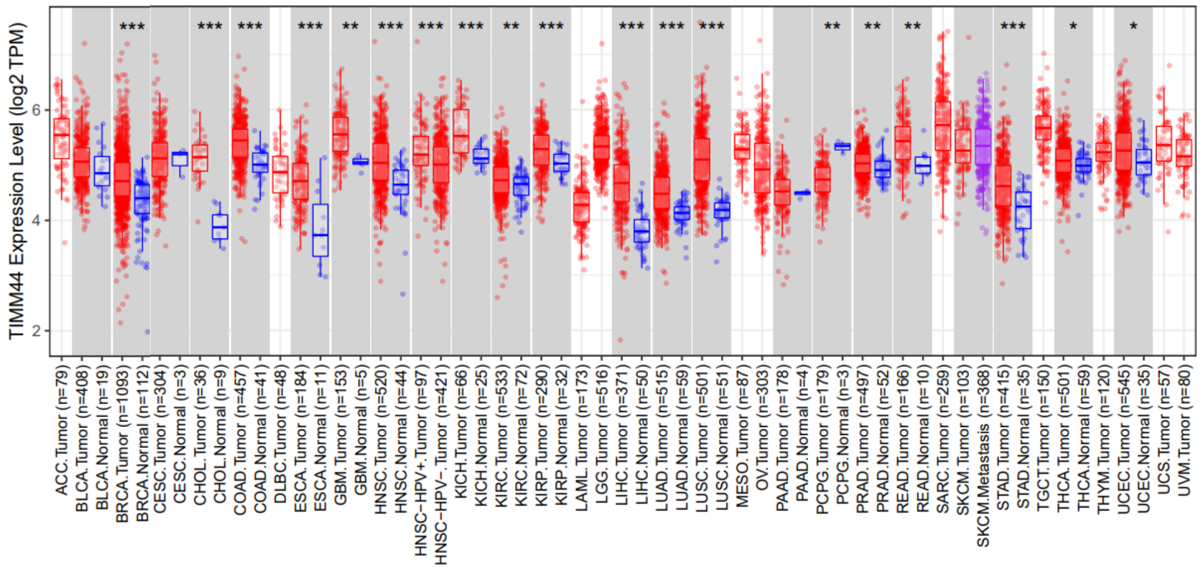


Immunohistochemical staining of Timm44 in normal and tumor tissues obtained from the Human Protein Atlas

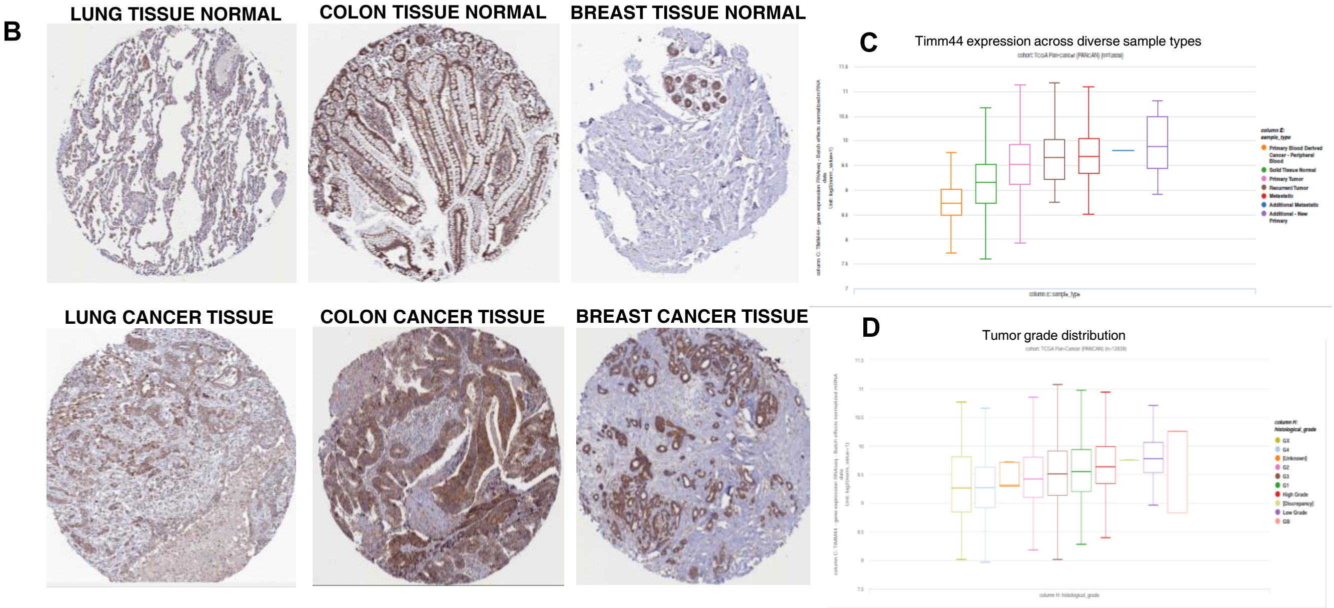

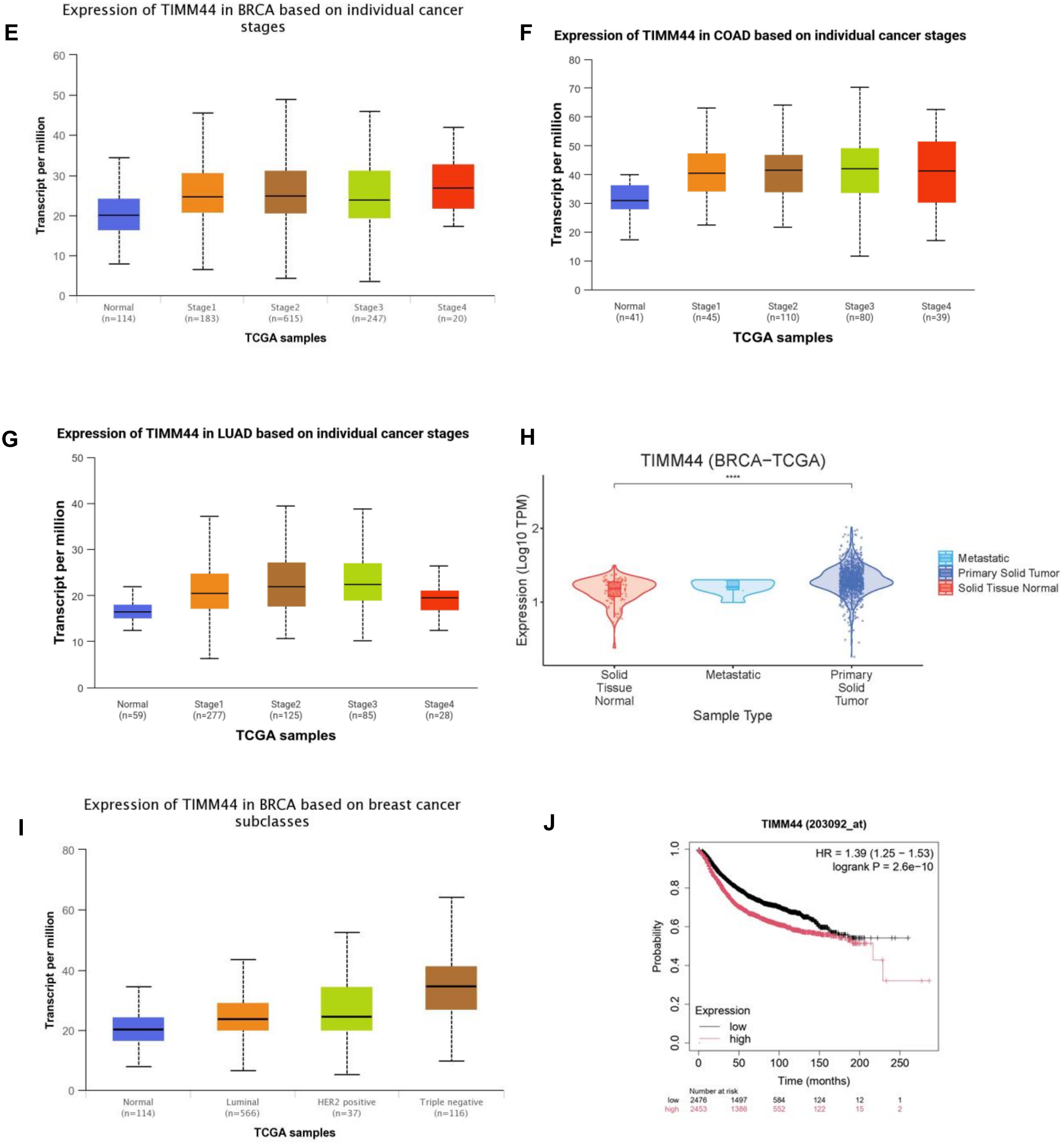
E
Expression of TIMM44 in BRCA based on individual cancer stages
Transcript per million
0
10
20
30
40
50
60
Normal (n=114)
Stage1 (n=183)
Stage2 (n=615)
Stage3 (n=247)
Stage4 (n=20)
TCGA samples
F
Expression of TIMM44 in COAD based on individual cancer stages
Transcript per million
0
10
20
30
40
50
60
70
80
Normal (n=41)
Stage1 (n=45)
Stage2 (n=110)
Stage3 (n=80)
Stage4 (n=39)
TCGA samples
G
Expression of TIMM44 in LUAD based on individual cancer stages
Transcript per million
0
10
20
30
40
50
Normal (n=59)
Stage1 (n=277)
Stage2 (n=125)
Stage3 (n=85)
Stage4 (n=28)
TCGA samples
H
TIMM44 (BRCA-TCGA)
****
Expression (Log10 TPM)
1
2
Solid Tissue Normal
Metastatic
Primary Solid Tumor
Sample Type
Metastatic
Primary Solid Tumor
Solid Tissue Normal
I
Expression of TIMM44 in BRCA based on breast cancer subclasses
Transcript per million
0
20
40
60
80
Normal (n=114)
Luminal (n=566)
HER2 positive (n=37)
Triple negative (n=116)
TCGA samples
J
TIMM44 (203092_at)
HR = 1.39 (1.25 − 1.53)
logrank P = 2.6e−10
Probability
0.0
0.2
0.4
0.6
0.8
1.0
Expression
low
high
0
50
100
150
200
250
Time (months)
Number at risk
low 2476 1497 584 124 12 1
high 2453 1386 552 122 15 2

K

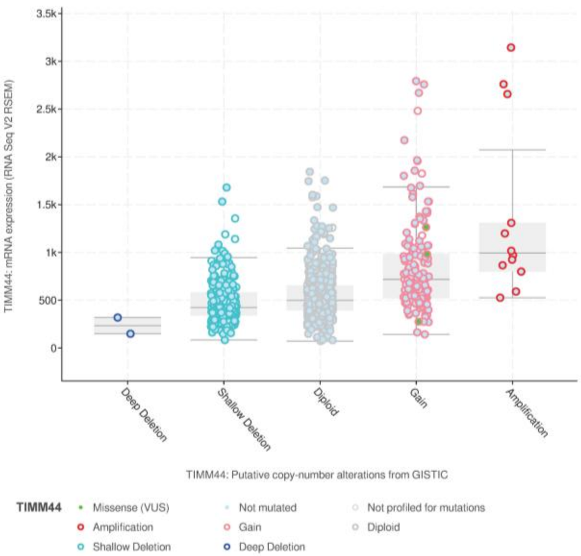


L

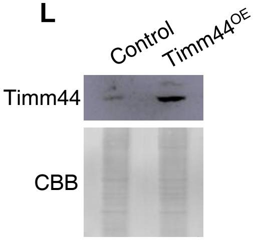


**Figure S1. Angiogenesis driven malignancies show elevated Timm44 expression**

**(A)** Expression of Timm44 across tumor and adjacent normal tissues in TCGA cancer types, obtained using the TIMER2 Gene_DE module. Gene expression distributions are shown as box plots, statistical significance was calculated using the edgeR-based differential expression analysis implemented in the TIMER2 and is indicated by asterisks. (**P* < 0.05; ***P* < 0.01; ****P* <0.001**). (B)** Immunohistochemical profiles of Timm44 protein expression obtained from the Human Protein Atlas (HPA). **(C)** Distribution of Timm44 expression across normal, primary tumor, and metastatic samples in the TCGA Pan-Cancer cohort, visualized using the UCSC Xena browser. Statistical significance was assessed by one-way ANOVA ($P = 0.000$, $F = 100.4$). **(D)** Association of Timm44 expression with tumor grade in TCGA Pan-Cancer cohort, analyzed using the UCSC Xena browser. Statistical significance was determined by one-way ANOVA ($P = 1.132 \times 10^{-14}$, $F = 9.616$) **(E-G)** Stage-wise expression analysis of Timm44 in Lung adenocarcinoma (LUAD), Colon adenocarcinoma (COAD), and Breast Cancer invasive carcinoma (BRCA) using UALCAN, with statistical significance assessed by pairwise Student's *t*-tests comparing normal tissue with each tumor stage. (*****P*<0.0001) **(H)**. Expression of TIMM44 across solid tissue normal, primary solid tumor, and metastatic samples in the TCGA-BRCA cohort (GEPIA2). Expression values are shown as $\log_2$ (TPM + 1), with statistical significance assessed using one-way ANOVA as implemented in GEPIA2. **(I)** Timm44 expression across molecular subclasses of breast cancer in the TCGA-BRCA dataset analyzed using UALCAN, with statistical significance assessed by pairwise Student's *t*-tests comparing normal tissue with each tumor stage. **(J)** Kaplan–Meier overall survival analysis based on Timm44 expression in the TCGA-BRCA cohort performed using the Kaplan–Meier Plotter. Survival differences were evaluated using the log-rank test. **(K)** Putative copy-number alteration profile of Timm44 in the TCGA Firehose Legacy breast cancer invasive carcinoma cohort, obtained from cBioPortal. (**L**) Immunoblot showing enhanced expression of Timm44 in MCF7 cells transfected with plasmid carrying Timm44 ORF.

### Figure S2

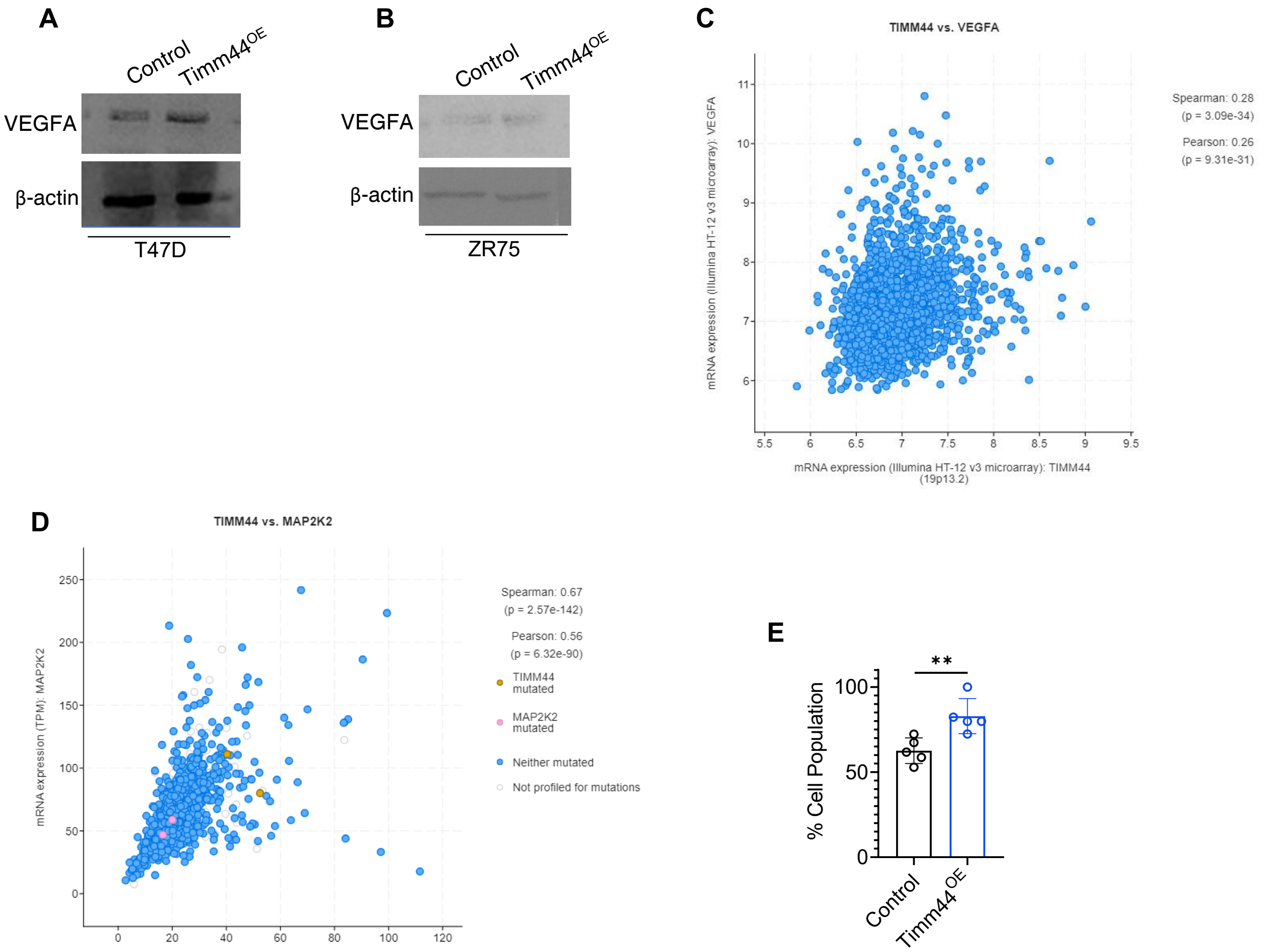
A
Control
Timm44$^{OE}$
VEGFA
β-actin
T47D
B
Control
Timm44$^{OE}$
VEGFA
β-actin
ZR75
C
TIMM44 vs. VEGFA
Spearman: 0.28
(p = 3.09e-34)
Pearson: 0.26
(p = 9.31e-31)
mRNA expression (Illumina HT-12 v3 microarray): VEGFA
mRNA expression (Illumina HT-12 v3 microarray): TIMM44 (19p13.2)
D
TIMM44 vs. MAP2K2
Spearman: 0.67
(p = 2.57e-142)
Pearson: 0.56
(p = 6.32e-90)
TIMM44 mutated
MAP2K2 mutated
Neither mutated
Not profiled for mutations
mRNA expression (TPM): MAP2K2
mRNA expression (TPM): TIMM44 (19p13.2)
E
**
% Cell Population
Control
Timm44$^{OE}$

F

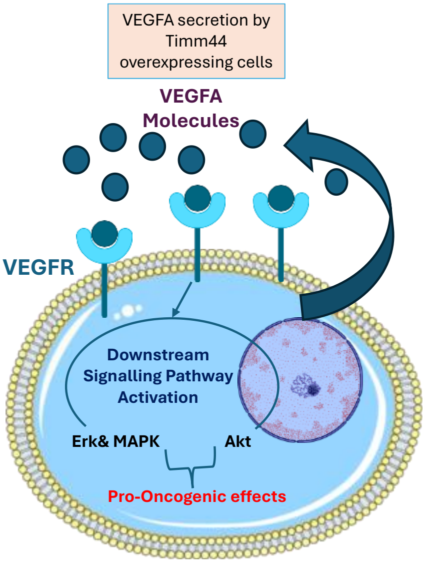


**Figure S2**

**(A, B)** Western blot analysis of VEGFA protein levels in control and Timm44$^{OE}$ T47D and ZR-75 breast cancer cell lines. **(C-D)** cBioPortal correlation analysis of Timm44 expression with VEGFA and MAP2K2 across TCGA breast cancer datasets. **(E)** MTT assay showing percent cell viability in control and Timm44$^{OE}$ cells. Data are presented as mean ± SEM (n = 5). Statistical significance was determined using Student's t-test, with ** indicating $p < 0.01$. **(F)** Schematic of VEGFA–VEGFR signaling and downstream ERK/MAPK-AKT pathways.

#### Figure S3

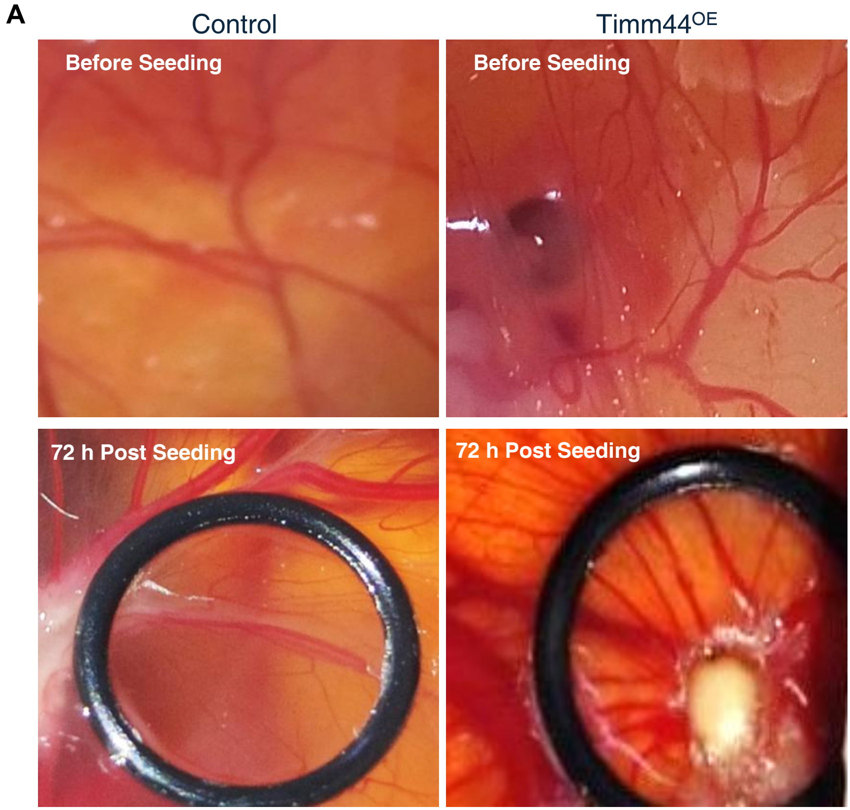


**Figure S3**
(A) 5000 control or Timm44-overexpressing cells were seeded on the CAM of 30 h chick embryo. Images were obtained 72 h post-seeding.

### Figure S4

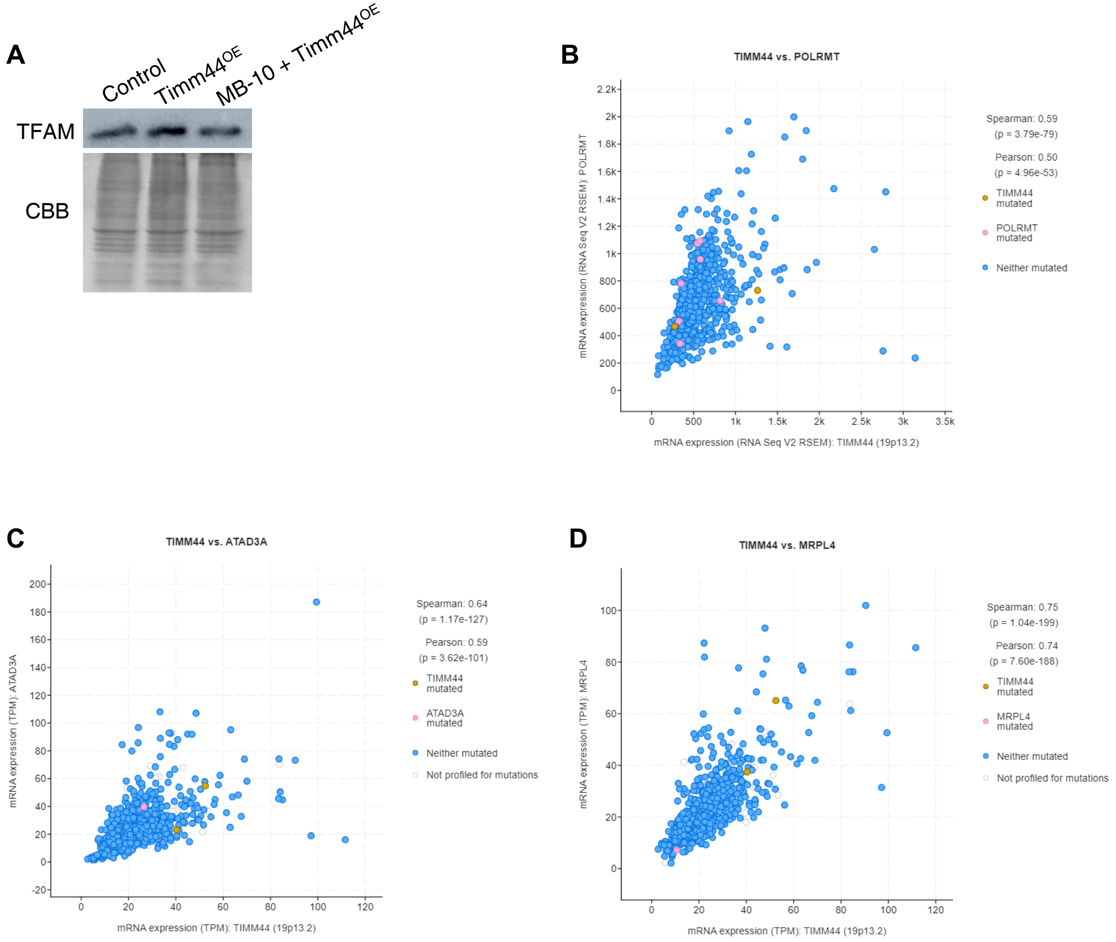

A
Control
Timm44OE
MB-10 + Timm44OE
TFAM
CBB
B
TIMM44 vs. POLRMT
mRNA expression (RNA Seq V2 RSEM): POLRMT
mRNA expression (RNA Seq V2 RSEM): TIMM44 (19p13.2)
Spearman: 0.59
(p = 3.79e-79)
Pearson: 0.50
(p = 4.96e-53)
TIMM44 mutated
POLRMT mutated
Neither mutated
C
TIMM44 vs. ATAD3A
mRNA expression (TPM): ATAD3A
mRNA expression (TPM): TIMM44 (19p13.2)
Spearman: 0.64
(p = 1.17e-127)
Pearson: 0.59
(p = 3.62e-101)
TIMM44 mutated
ATAD3A mutated
Neither mutated
Not profiled for mutations
D
TIMM44 vs. MRPL4
mRNA expression (TPM): MRPL4
mRNA expression (TPM): TIMM44 (19p13.2)
Spearman: 0.75
(p = 1.04e-199)
Pearson: 0.74
(p = 7.60e-188)
TIMM44 mutated
MRPL4 mutated
Neither mutated
Not profiled for mutations

E

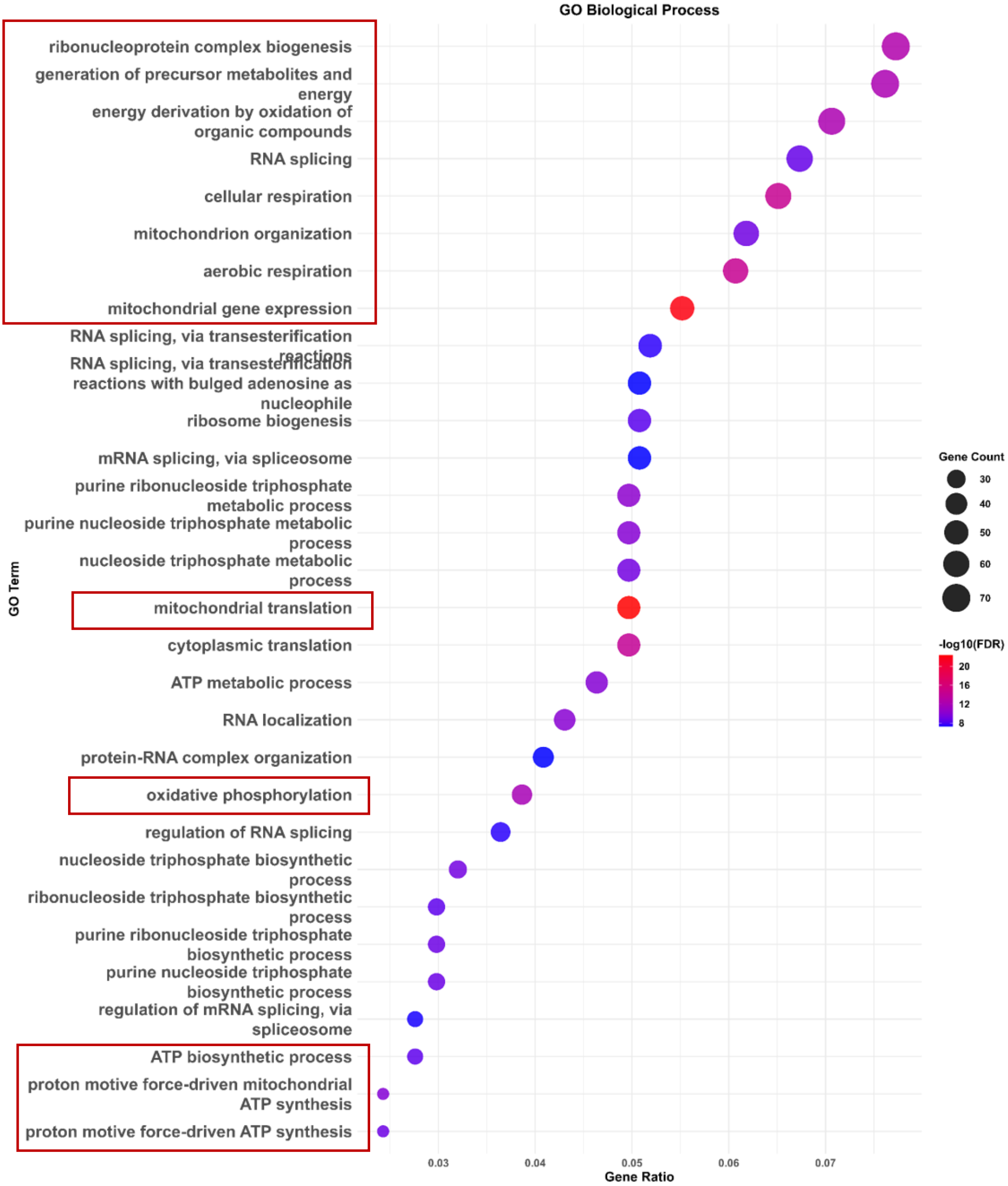
GO Biological Process
ribonucleoprotein complex biogenesis
generation of precursor metabolites and energy
energy derivation by oxidation of organic compounds
RNA splicing
cellular respiration
mitochondrion organization
aerobic respiration
mitochondrial gene expression
RNA splicing, via transesterification reactions
RNA splicing, via transesterification reactions with bulged adenosine as nucleophile
ribosome biogenesis
mRNA splicing, via spliceosome
purine ribonucleoside triphosphate metabolic process
purine nucleoside triphosphate metabolic process
nucleoside triphosphate metabolic process
mitochondrial translation
cytoplasmic translation
ATP metabolic process
RNA localization
protein-RNA complex organization
oxidative phosphorylation
regulation of RNA splicing
nucleoside triphosphate biosynthetic process
ribonucleoside triphosphate biosynthetic process
purine ribonucleoside triphosphate biosynthetic process
purine nucleoside triphosphate biosynthetic process
regulation of mRNA splicing, via spliceosome
ATP biosynthetic process
proton motive force-driven mitochondrial ATP synthesis
proton motive force-driven ATP synthesis
GO Term
0.03
0.04
0.05
0.06
0.07
Gene Ratio
Gene Count
30
40
50
60
70
-log10(FDR)
20
16
12
8

**Figure S4**

**(A)** Western blot analysis of TFAM protein levels in control, $Timm44^{OE}$, and $Timm44^{OE}$ cells treated with MB-10, further quantified by densitometry and represented as fold change over control. **(B-D)** cBioPortal correlation analysis of Timm44 expression with POLRMT, ATAD3, and MRPL4. **(E)** GO biological process enrichment of upregulated and downregulated genes in $Timm44^{OE}$ cells relative to control, assessed by ORA. $-\log_{10}(FDR)$ values are shown by color scale, and dot size reflects gene counts.

### Figure S5

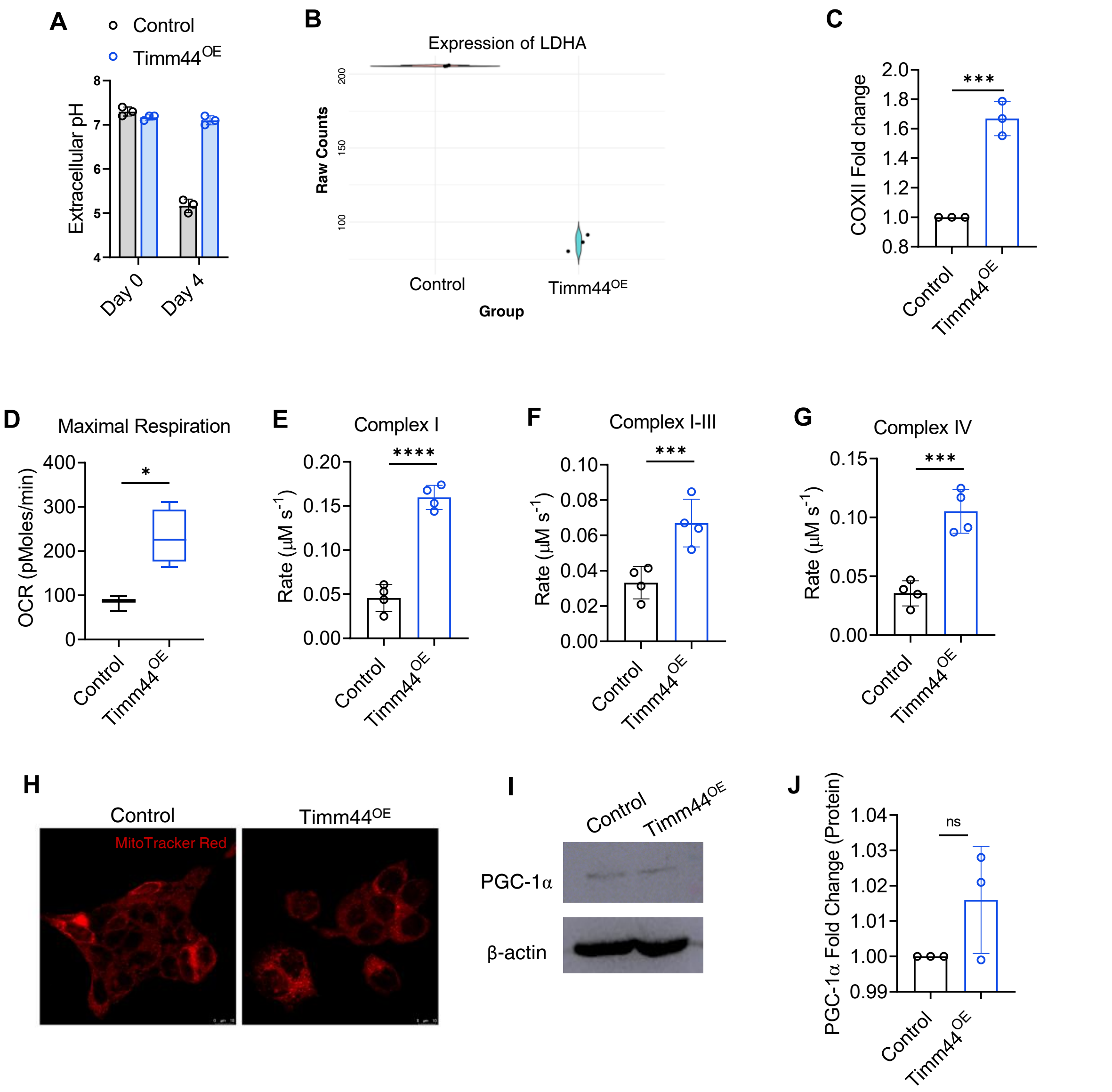
A
Control
Timm44OE
Extracellular pH
Day 0
Day 4
B
Expression of LDHA
Raw Counts
Group
Control
Timm44OE
C
COXII Fold change
***
Control
Timm44OE
D
Maximal Respiration
OCR (pMoles/min)
*
Control
Timm44OE
E
Complex I
Rate (μM s-1)
****
Control
Timm44OE
F
Complex I-III
Rate (μM s-1)
***
Control
Timm44OE
G
Complex IV
Rate (μM s-1)
***
Control
Timm44OE
H
Control
Timm44OE
MitoTracker Red
I
Control
Timm44OE
PGC-1α
β-actin
J
PGC-1α Fold Change (Protein)
ns
Control
Timm44OE

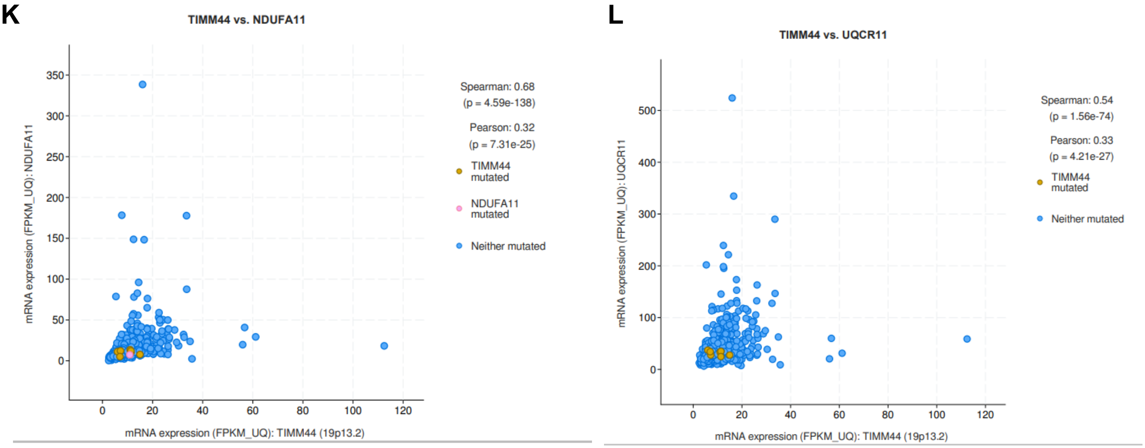


**Figure S5**

(**A**) Changes in extracellular pH caused by control and Timm44-overexpressing cells, measured on day 0 and day 4, n = 3. **(B)** LDHA protein abundance derived from raw proteomics counts. **(C)** COXII protein fold change determined by densitometric analysis of Fig 4A COXII blot, represented as fold change over control, n = 3. **(D)** Maximal respiration measured by oxygen consumption rate (OCR), n = 3 **(E-G)** Assessment of Complex I, Complex IV and Complex I-III activities using isolated mitochondria derived from Timm44 overexpressing and control cells. n =4. **(H)** Mitochondrial morphology assessed by MitoTracker staining. **(I-J)** PGC-1α protein levels assessed by immunoblotting, quantified by densitometric analysis and represented as fold change over control, n = 3. All data are presented as mean ± SEM. Statistical significance was determined using two-tailed Student's t-tests. $*P < 0.05$, $**P < 0.01$, $***P < 0.001$, $**P < 0.0001$. **(K-L)** cBioPortal correlation analysis of Timm44 expression with NDUFA11 (Complex I subunit) and UQCR11 (Complex III subunit).

### Figure S6

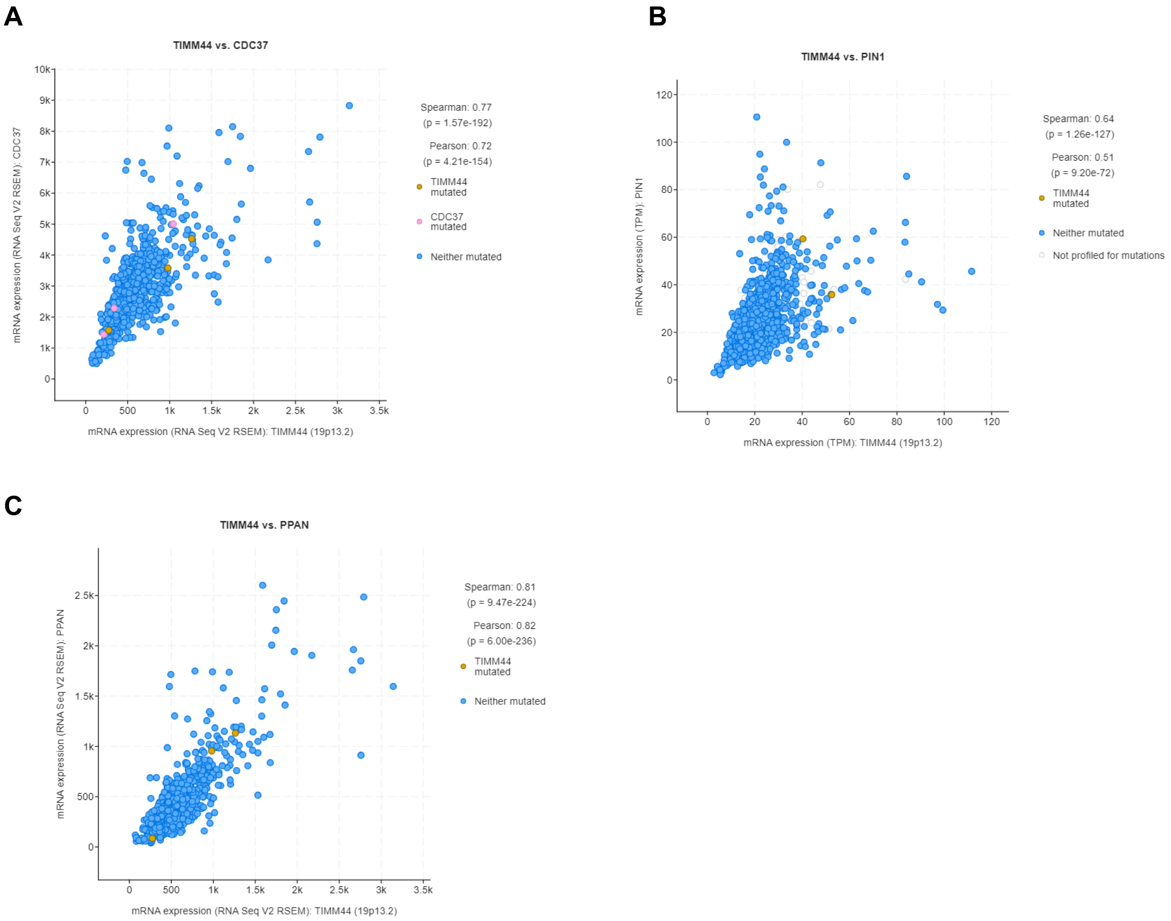


**Figure S6**
**(A-D)** cBioPortal correlation analyses showing associations between Timm44 expression and the regulatory factors CDC37, PIN1 and PPAN.

**Figure S7**

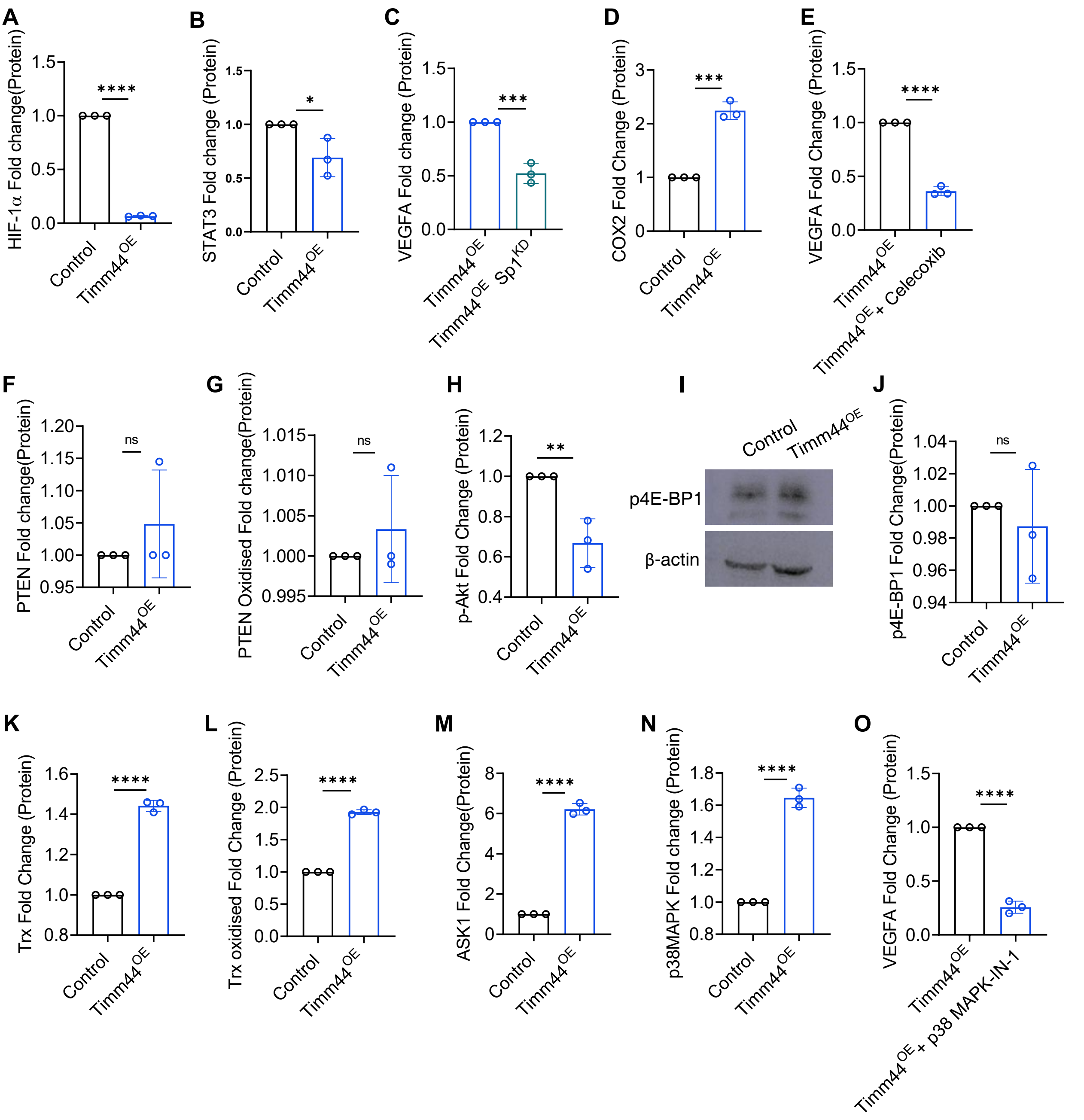
A
HIF-1α Fold change(Protein)
Control
Timm44OE
****
B
STAT3 Fold change (Protein)
Control
Timm44OE
*
C
VEGFA Fold change (Protein)
Timm44OE
Timm44OE Sp1KD
***
D
COX2 Fold Change (Protein)
Control
Timm44OE
***
E
VEGFA Fold Change (Protein)
Timm44OE
Timm44OE + Celecoxib
****
F
PTEN Fold change(Protein)
Control
Timm44OE
ns
G
PTEN Oxidised Fold change(Protein)
Control
Timm44OE
ns
H
p-Akt Fold Change (Protein)
Control
Timm44OE
**
I
Control
Timm44OE
p4E-BP1
β-actin
J
p4E-BP1 Fold Change(Protein)
Control
Timm44OE
ns
K
Trx Fold Change (Protein)
Control
Timm44OE
****
L
Trx oxidised Fold Change (Protein)
Control
Timm44OE
****
M
ASK1 Fold Change(Protein)
Control
Timm44OE
****
N
p38MAPK Fold change (Protein)
Control
Timm44OE
****
O
VEGFA Fold Change (Protein)
Timm44OE
Timm44OE + p38 MAPK-IN-1
****

**Figure S7**

**(A, B)** Densitometric quantification of Western blots for HIF1α and STAT3 in Fig 6B,C; represented as fold change over control, n = 3. **(C)** Quantification of VEGFA levels upon SP1 knock-down, as in Fig. 6F, represented as fold change over Timm44$^{OE}$ cells, n=3. **(D)** Quantification of Cox2 levels in Fig 6G represented as fold change over controls, n=3. **(E)** Quantification of VEGFA levels following COX-2 inhibition by celecoxib inhibitor, Fig 6H, represented as fold change over control, n = 3. **(F-G)** Densitometric quantification of Western blots for PTEN and oxidized PTEN (Fig 6J,K), represented as fold change over control, n = 3. **(H)** Densitometric quantification of Western blots for pAkt in Fig 6L; represented as fold change over control, n = 3. **(I, J)** Western blot analysis of p-4EBP protein levels quantified as fold change over control (J), n = 3. **(K-N)** Densitometric quantification of Western blots for TRX and oxidized TRX, ASK, p38 MAPK (Fig 6M-P), n=3. **(O)** Quantification of VEGFA levels following p38 inhibition by p38MAPK-IN-1 inhibitor, represented as fold change over control (Fig 6R), n = 3. All data are presented as mean ± SEM. Comparisons were performed between control and Timm44$^{OE}$ cells, or Timm44$^{OE}$ cells and Timm44$^{OE}$ cells transfected with SP1 siRNA/treated with an inhibitor (p38 MAPK-IN-1 or celecoxib). Statistical significance was determined using two-tailed Student's t-tests. $P < 0.05$, $P < 0.01$, $P < 0.001$, $P < 0.0001$.
